# Multi-omic Analysis of Primary Human Kidney Tissues Identifies Medulla-Specific Gene Expression Patterns

**DOI:** 10.1101/2022.10.05.508277

**Authors:** Stefan Haug, Selvaraj Muthusamy, Yong Li, Anna Köttgen, Shreeram Akilesh

**Affiliations:** Institute of Genetic Epidemiology, Medical Center-University of Freiburg, Freiburg, Germany; Department of Pathology, Virginia Commonwealth University, Richmond, USA; Department of Laboratory Medicine and Pathology, University of Washington, Seattle, USA

## Abstract

The renal medulla is a specialized region of the kidney with important homeostatic functions. It has also been implicated in genetic and developmental disorders and ischemic and drug-induced injuries. Despite its role in kidney function and disease, the medulla’s baseline gene expression and epigenomic signatures have not been well described in the adult human kidney. Here we generate and analyze gene expression (RNA-seq), chromatin accessibility (ATAC-seq) and chromatin conformation (Hi-C) data from adult human kidney cortex and medulla. Using data from our carefully annotated specimens, we assign samples in the larger public GTEx database to cortex and medulla, thereby identifying several misassignments and extracting meaningful medullary gene expression signatures. Using integrated analysis of gene expression, chromatin accessibility and conformation profiles, we reveal insights into medulla development and function. Our datasets will also provide a valuable resource for researchers in the GWAS community for functional annotation of genetic variants.

## Introduction

The kidney medulla has important homeostatic functions as it contains the portions of the nephron that fine tune salt and water balance in the circulation. It is also a target of genetic kidney diseases such as autosomal-dominant tubulointerstitial kidney disease (ADTKD-UMOD) and certain congenital abnormalities of the renal and urinary tract. The kidney medulla has a specialized hypoxic and high salt milieu ^1,2^ and represents a specialized immune environment that resists infection ^3^. The medulla is particularly sensitive to ischemic injury and certain nephrotoxins. Despite these important roles, the expression patterns and regulation of genes important to human kidney medulla function are relatively understudied. The large Genotype-Tissue Expression (GTEx) project ^4^, which is widely used for the study of tissue-specific gene expression, contains 85 samples of kidney cortex, but only 4 samples of kidney medulla. Another widely used resource, the Human Protein Atlas ^5^, does not contain any expression data for human kidney medulla. Furthermore, epigenomic data such as chromatin accessibility and conformation, are not available for the kidney medulla. Generation of such data sets from primary tissue samples is important since cultured cells alter their epigenomic features and can show an injured phenotype ^6^.

Together, these deficits have hampered our understanding of the medulla’s role in numerous kidney diseases, including developmental defects but also acute kidney injury. To address this knowledge gap, here we generate human kidney cortex- and medulla-specific gene expression (RNA-seq), chromatin accessibility (ATAC-seq) and genome organization (Hi-C) datasets. Through integrative and comparative analyses between the cortex and medulla, we provide new insights into medulla-specific gene expression and regulation. We demonstrate an approach of using a small set of carefully curated human tissue samples to detect and correct for tissue-misassignment in publicly available gene expression data, which has implications for the use of GTEx tissue samples beyond the kidney. The medulla-specific datasets resulting from our study represent a resource for experimental researchers working on medullary genes as well as the GWAS community for functional annotation.

## Results

### Differential gene expression between kidney cortex and medulla

We performed RNA-seq and ATAC-seq on kidney tissue from three male human kidney donors, which was each macrodissected into cortical and medullary tissue samples (Fig. 1). Histologic evaluation at the timepoint of sampling revealed good representation of the respective kidney region (Fig. 2a). All three patients (age 57, 67 and 73) had preserved kidney function (eGFR >70 ml/min/1.73m^2^; Fig. 2a, Supplementary Table 1), and histologic review by a renal pathologist (S.A.) showed only mild arteriosclerosis and <5% nephrosclerosis (global glomerulosclerosis, tubular atrophy and interstitial fibrosis). Histological assessment also confirmed accurate macrodissection into cortex or medulla (Fig. 2b, Supplementary Fig. 1). Consistent with the histology results, principal component analysis (PCA) based upon both gene expression (RNA-seq) and chromatin accessibility (ATAC-seq) data showed clear separation between cortical and medullary samples (Fig. 2c). Using snRNA-seq data from the KPMP Kidney Tissue Atlas ^7^ as a reference, we deconvoluted the estimated abundance of different kidney cell types in our bulk RNA-seq data using BisqueRNA ^8^ (Figure 2d; Supplementary Table 2). Reassuringly, the most significant differences were found for proximal tubule cells, which were enriched in our cortical samples (42% +/- 3% compared to 22% +/- 5 % in medulla, p-value 0.002), and cells of the thick ascending limb which were more abundant in our medullary samples (37% +/- 4% compared to 17% +/- 2% in cortex, p-value 0.004). These histologic and computational assessments therefore validated the correct assignments of cortical and medullary tissue samples for subsequent comparative studies.

**Figure 1:**
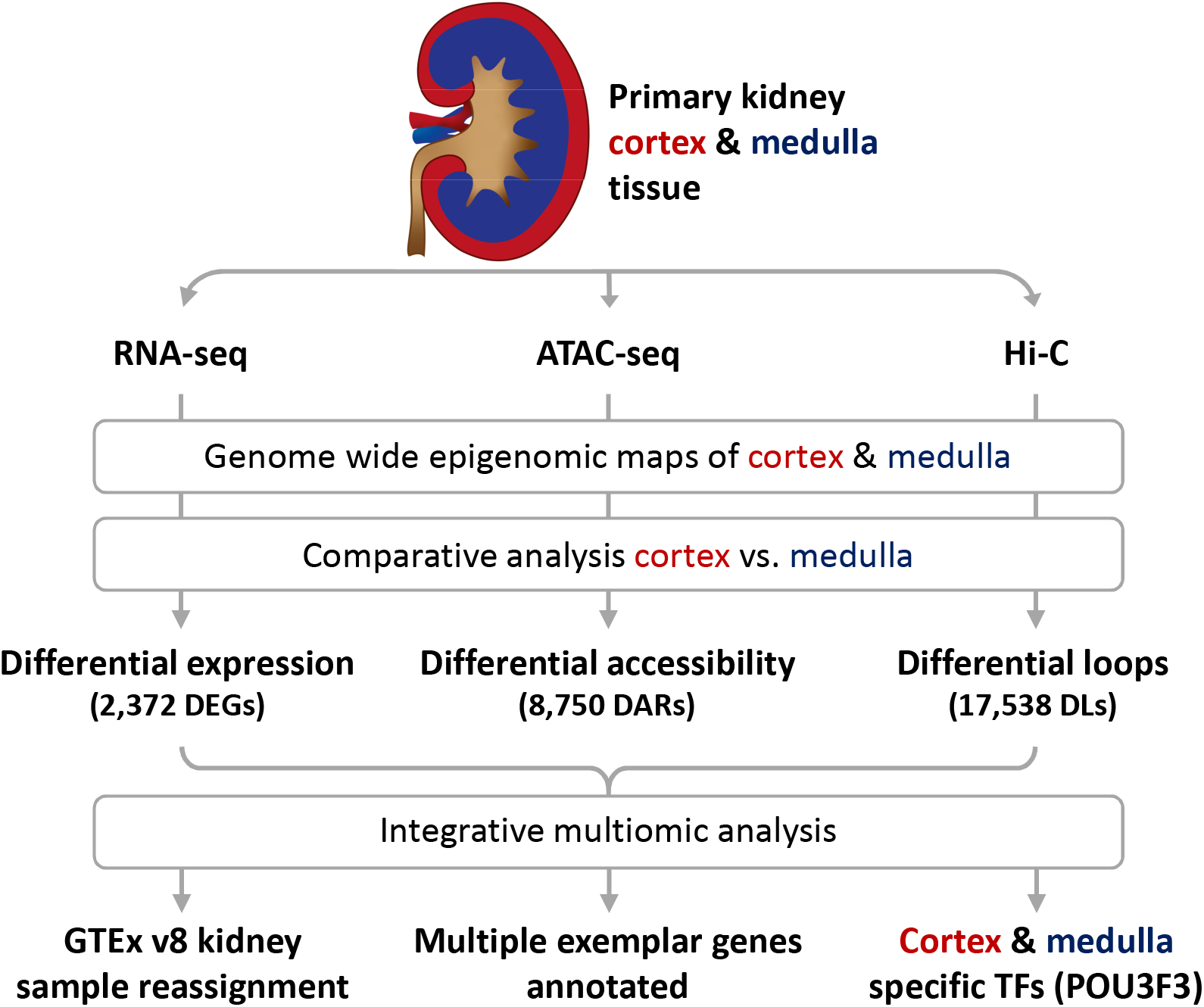
Overview of sample types and analysis workflow. A flowchart depicting sample processing and analysis workflows utilized in this manuscript.

**Figure 2:**
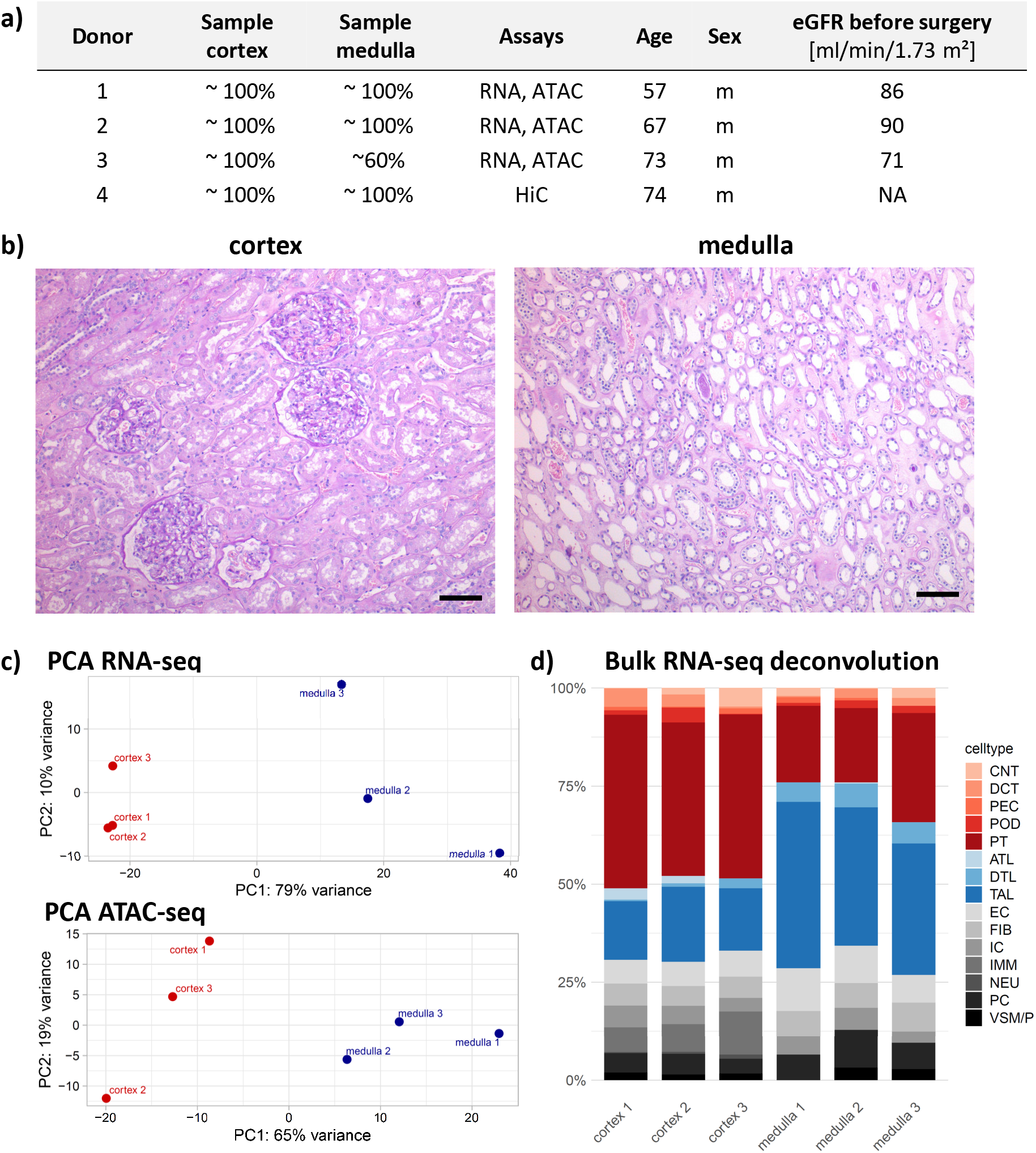
Composition of tissue samples. **a)** Histological assessment of sample purity, assays performed with tissue samples and clinical characteristics of tissue donors (additional information in Supplementary Table 1). The estimated glomerular filtration rate (eGFR) was computed with the CKD-EPI creatinine equation (2021) ^63^. NA, not available. **b)** Representative sections of macrodissected cortex and medulla tissue from donor 1 (PAS staining). Scale bar, 100µm. **c)** PCA of all 6 samples from donor 1 – 3 based on RNA-seq data (top, using counts per transcript) and ATAC-seq data (bottom, using counts per accessible region). PC1 explains the largest proportion of variance between samples in both assays (79% for RNA-seq and 65% for ATAC-seq). **d)** Deconvolution analysis of bulk RNA-seq data with BisqueRNA showing the estimated proportions of different cell types in our tissue samples. Cortical cell types are shaded in red, medullary cell types in blue, cell types without specified localization are shown in grey. Cell type abbreviations include podocytes (POD), parietal epithelial cells (PEC), descending and ascending thin limb cells (DTL, ATL), proximal tubules (PT), thick ascending limb (TAL), distal convoluted tubule cells (DCT), connecting tubule cells (CNT), epithelial cells (EC), fibroblasts (FIB), intercalated cells (IC), parietal cells (PC), immune cells (IMM), neuronal cells (NEU), and vascular smooth muscle cells / pericytes (VSM/P).

To begin, we performed a differential gene expression analysis between the cortical and medullary samples with DESeq2 ^9^ (Methods). Among 25,185 genes with at least minimal detectability (>= 5 detected transcripts in >= 2 samples), we found 2,372 or 9.4% to be differentially expressed between cortex and medulla (adjusted p-value < 0.01, log_2_-fold change > 1; Supplementary Table 3). There were 1,266 differentially expressed genes (DEGs) with higher expression in cortex and 1,106 DEGs with higher expression in medulla (Supplementary Fig. 2a). Next, we ranked DEGs based on their expression differences both in terms of absolute counts as well as in terms of statistical significance (i.e., adjusted p-value) in order to identify genes with the strongest support for between-group differences. Reassuringly, this revealed genes with highly differential expression in the medulla, including the canonical genes *UMOD, SLC12A1* and *AQP2* (Supplementary Fig. 2b, c). In gene ontology over-representation analysis, while cortical DEGs were enriched for transporter activity and metabolic processes, medullary DEGs were enriched for terms related to angiogenesis, extracellular matrix and renal or urogenital system development (Supplementary Fig. 3; Supplementary Tables 4 and 5).

### Carefully annotated samples recalibrate and extend the utility of the larger GTEx dataset

Next, we asked if we could replicate the detected expression differences in bulk kidney RNA-seq data for 85 cortical and 4 medullary samples generated by the GTEx project. Surprisingly, performing DEG analysis on these GTEx cortex and medulla samples yielded only a very low number of 76 DEGs (Fig. 3a) without any significantly enriched GO-terms. To investigate this apparent discrepancy, we first performed a principal component analysis (PCA) based on the GTEx data (Methods). This showed a separation of the samples into two clusters along principal components (PC) 1 and 2, the smaller of which contained 9 samples. However, the clusters did not clearly correspond to the GTEx sample assignment to cortex or medulla (Fig. 3c), as we had observed in the PCA of our own carefully curated samples. When we examined the published pathology descriptors of the GTEx kidney samples, we noted that several samples in the smaller GTEx cluster described contamination from the opposite category. These findings supported a potential mis-assignment of cortex samples as medulla and vice versa.

**Figure 3.**
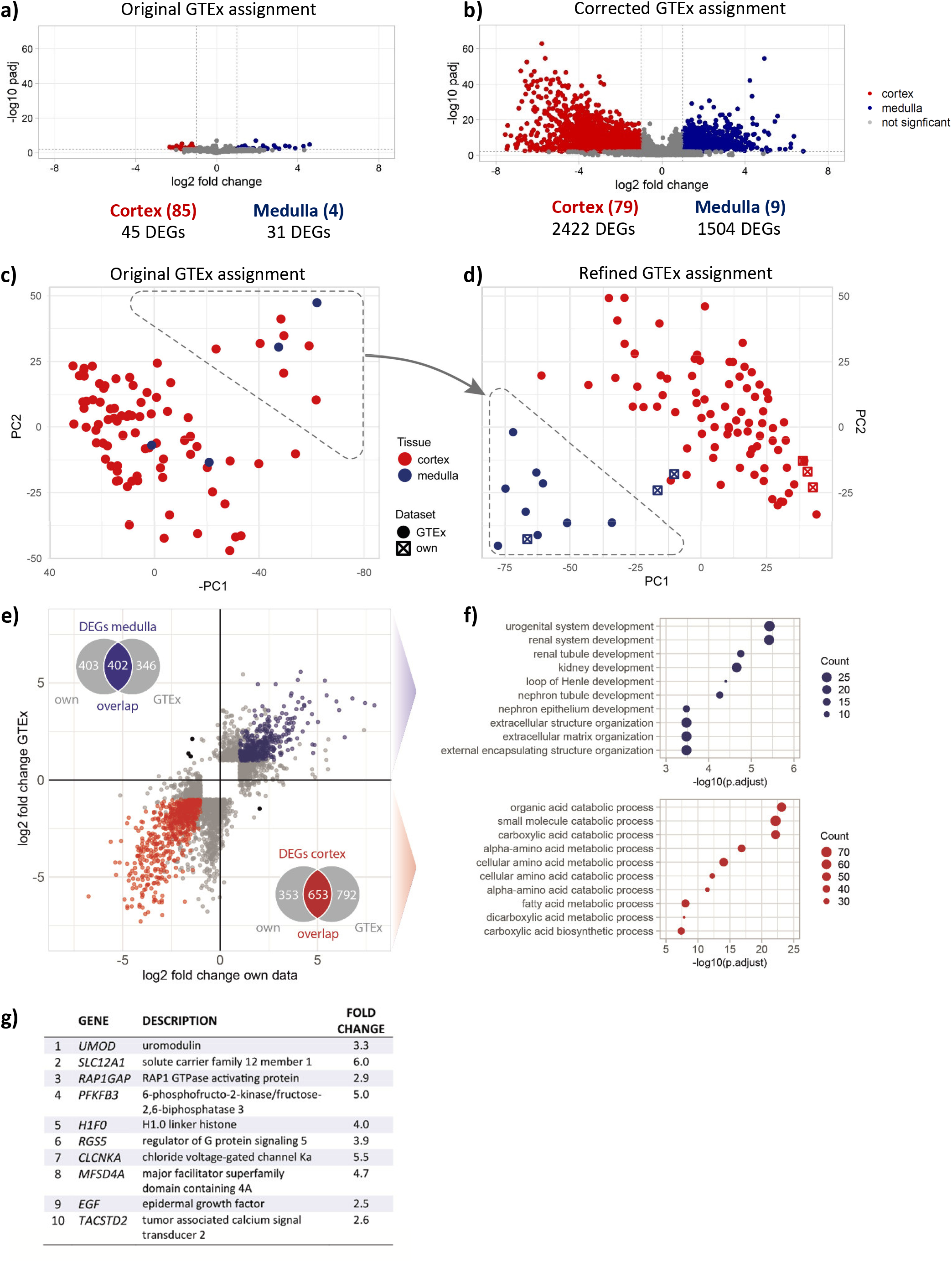
GTEx v8 kidney sample re-calibration and differential expression analysis. **a)** Volcano plot and number of DEGs from the differential gene expression analysis comparing cortex and medulla samples in the GTEx V8 database (significance thresholds indicated with dotted lines: log2 fold change > 1, padj < 0.01). Fold changes were computed as medulla/cortex, i.e., positive log2 fold changes mean higher expression in medulla, negative values higher expression in cortex. **b)** Volcano plot and number of DEGs after correction of sample assignment shown with the same axes as in panel a. **c)** The PCA based on GTEx RNA-seq data. Color coding highlights the original assignment of samples to cortex and medulla. Note the mixed tissue types in both sample clusters. **d)** Combined PCA of our own and the GTEx samples. Color coding highlights the corrected assignment to cortex and medulla for the GTEx samples. GTEx samples from the top right cluster in panel c are now located close to our medullary samples. The remaining GTEx samples lie closer to our cortical samples. **e)** Overlap of DEGs from our own and the corrected GTEx samples. 402 medullary and 653 cortical DEGs were identified in both analyses. Overlapping DEGs show highly correlating log2 fold changes (Spearman’s ρ = 0.92, red and blue dots in scatterplot), only 4 DEGs (<1%) show log2 fold changes of opposing directions (black dots). **f)** Enriched GO-terms (molecular function) for overlapping DEGs (top: medulla, bottom: cortex) with adjusted p-value for enrichment and number of genes belonging to the respective terms (count). **g)** DEGs identified from our and the corrected GTEx samples with the highest average expression in medulla across both datasets (mean of normalized counts from both datasets).

We therefore adopted a PCA-based approach widely used in genetic epidemiology for the assignment of genetic ancestry of samples of unknown ancestry, when a reference dataset of known ancestries is available ^10,11^. Using our own cortex and medulla samples as reference, we performed a joint PCA analysis with the GTEx samples, which is agnostic to the assignment of the tissue samples in GTEx. Interestingly, we observed that the exact same 9 GTEx samples detected as a smaller cluster of the GTEx-only PCA clustered together with our medullary samples (Fig. 3d), supporting potential misassignment of several GTEx medulla samples as cortex. Indeed, 6 of these 9 GTEx samples had pathology information available that described contamination (Supplementary Fig. 4). Therefore, we reassigned the GTEx samples to medulla and cortex based on the clustering observed in the joint PCA analysis (Fig. 3d) as described in the Methods, yielding 9 re-assigned GTEx medulla and 80 GTEx cortex samples for downstream analyses (Supplementary Table 6). After reassignment, the number of GTEx DEGs increased sharply from 76 to 3,926 (Fig. 3 b). Correct re-assignment was supported by over-representation of similar gene ontology terms as detected in our own reference samples.

Next, we intersected the resulting GTEx DEGs with the 2,372 DEGs from our own samples. We identified 1,055 overlapping, significant DEGs (653 with higher expression in the cortex, 402 with higher expression in the medulla; Supplementary Table 8). These overlapping DEGs exhibited highly correlated log_2_-fold changes across the two datasets (Spearman’s ρ = 0.92, Fig. 3e), which was even higher than the correlation for all DEGs alone (Spearman’s ρ = 0.75). The DEGs identified across both datasets were biologically plausible: First, medullary DEGs with high average expression included genes with well-established roles in diseases originating in the medulla such as *UMOD* and *SLC12A1* (Fig. 3g). Second, overlapping medullary DEGs were overrepresented in Gene Ontology (GO)-terms related to urogenital system or kidney development and the organization of extracellular matrix (Fig. 3f). Conversely, cortical DEGs included members of the mitochondrial cytochrome c oxidase complex (*MT-CO1, MT-CO3*) and metabolic enzymes such as aldolase B (*ALDOB*), which is consistent with the high metabolic activity and energy demand of proximal tubule epithelial cells in the cortex. In addition, many solute transporters expressed in proximal and distal tubular cells that are over-represented in the cortex were highly expressed, for example *SLC12A3*, the renal thiazide-sensitive sodium-chloride cotransporter that is a marker gene for the distal convoluted tubule. The overlapping cortical DEGs were overrepresented for terms related to metabolic processes of small molecules handled by the kidney’s tubular cells (Fig. 3f). Therefore, using only a small number of carefully annotated samples, we were able to recalibrate and extend the utility of larger public datasets such as GTEx.

### Integrative analysis of gene expression, chromatin accessibility and conformation data

We subsequently proceeded to analyze the chromatin accessibility (ATAC-seq) data that we had generated on these same samples. We first identified accessible chromatin regions (ARs) in all cortical and medullary samples and then merged these regions into a master list yielding a total of 196,278 ARs (Methods). Of these, 8,750 (4.5%) showed differential accessibility between cortex and medulla and were designated as differentially accessible regions (DARs). Of the DARs, 5,462 had higher accessibility in cortex, while 3,288 displayed higher accessibility in the medulla (Fig. 4a).

**Figure 4:**
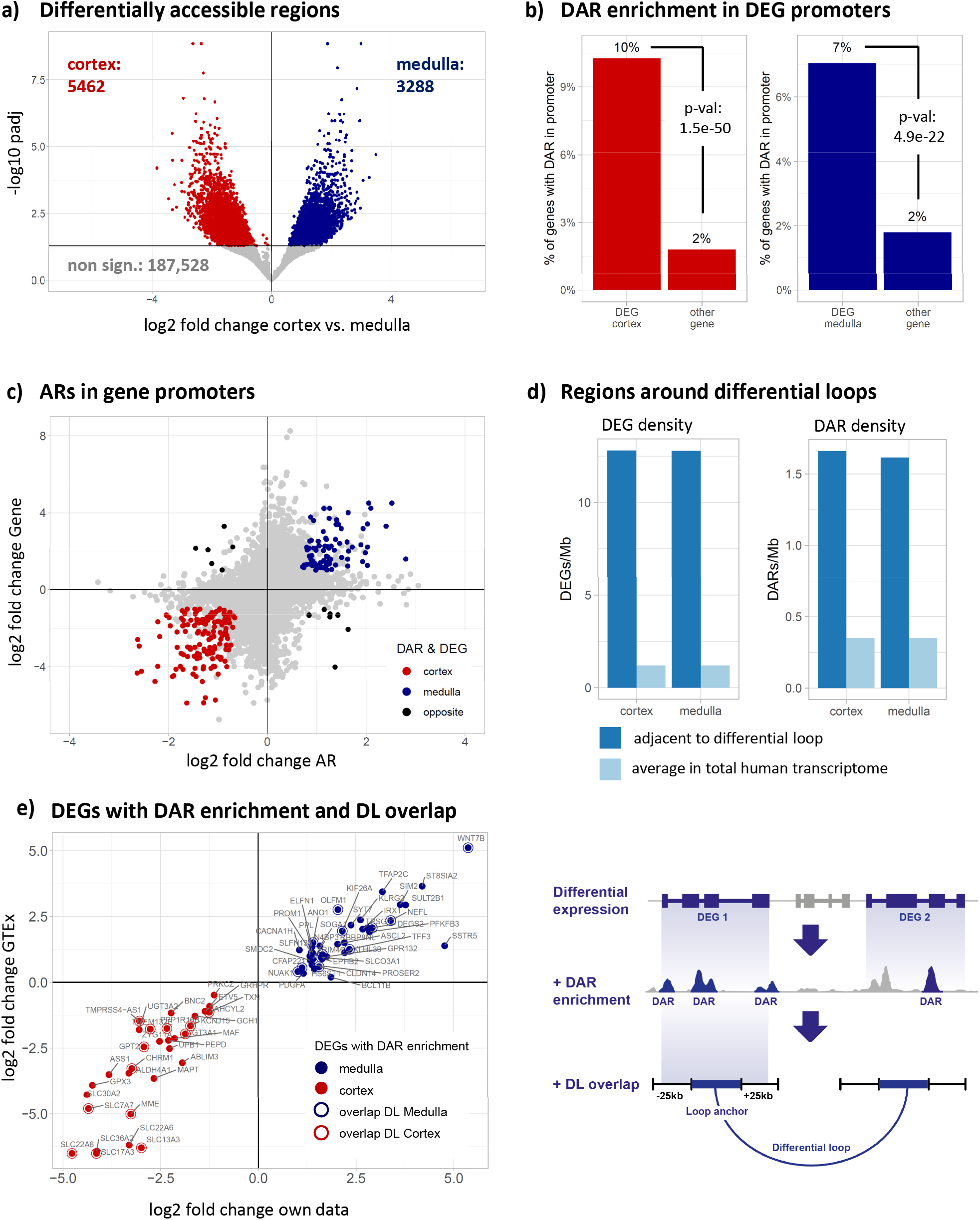
Differential analysis of cortex and medulla with RNA-seq, ATAC-seq and Hi-C data. **a)** Comparing the accessibility at 196,278 ARs in our tissue samples yielded 5,462 cortical and 3,288 medullary DARs (padj < 0.05, indicated by horizontal line). **b)** Percentage of genes (DEGs and non-DEGs) which have DARs in their promoter region (−5000 bp to 1000 bp from TSS). DEGs have a significantly higher proportion of genes with DARs in their promoters (cortex: 130/1,266, medulla: 78/1106) than non DEGs (408/22,753). P-values from Fisher’s exact test. **c)** Correlation of log2 fold change in accessibility medulla vs. cortex with the log2 fold change in gene expression for all ARs located in gene promoters (−5000 bp – 1000 bp from TSS). Colors: DARs located in DEG promoters (r = 0.82), grey: All other accessible regions in promoters (r = 0.19). **d)** Density of DEGs and DARs in genomic regions close to differential chromatin loops. The 10 kb loop anchors were extended by 25 kb to both sides. Then the density of DEGs and DARs per Mb was computed for these regions and compared to the average density of DARs and DEGs in the total human transcriptome (background). **e)** DEGs from our own samples were screened for DAR enrichment in their extended regulatory region and then overlapped with the loop anchors of DLs (see schematic illustration to the right and Methods). We identified 71 DEGs with DAR enrichment, all of which showed consistent log2 fold changes in the GTEx data (r = 0.94). 25 genes additionally overlapped extended regions around DL anchors.

Most ARs and DARs were found in promoter regions, exons and introns (71-80%), with the remaining ARs located in downstream and intergenic regions (Supplementary Fig. 5, Supplementary Table 8). DEGs were much more likely than non-differentially expressed genes to have a DAR within their promoter (defined as −5kB to +1kB from TSS, Methods). For example, 10% (130/1,266) of cortical and 7% (78/1106) of medullary DEGs had at least one DAR within their promoter compared to 2% (408/22,753) of all non-DEGs (Fig. 4b). Furthermore, DARs in proximity to DEGs showed a strong positive correlation between changes in accessibility and the DEG’s expression (Fig. 4c; Supplementary Fig. 6). The correlation was most prominent for DARs located in promoter regions (r = 0.82) and directly downstream of the gene body (r = 0.85). This correlation was less pronounced for non-differential ARs and genes (r from 0.09 to 0.25 depending on AR location, Supplementary Table 12). We then identified the subset of DEGs with an enrichment of DARs in a broader regulatory region around genes including the promoter, the gene body and additional up- and downstream regions. We intersected these DEGs with those genes also showing significant expression differences in the GTEx data. This identified 39 medullary and 32 cortical DEGs with DAR enrichment (Supplementary Table 13), which represents the gene set with the highest level of concordance across gene expression and chromatin accessibility in both our data and in GTEx (Fig. 4e). As for the DEGs, many of the DEGs with DAR enrichment are plausible biological candidates for the compartment in which they were detected, as exemplified by several solute carriers involved in transmembrane transport processes of proximal tubular cells that were detected in cortex.

Accessible chromatin elements such as enhancers can make long-range (multi-kilobase) interactions with their target genes. To study such chromatin contacts, we generated high-resolution genome-wide chromatin conformation data (Hi-C) from one matched cortex-medulla pair. We mapped 369,485,049 contacts in the cortex and 423,001,336 contacts in the medulla sample, the majority of which were short-range. In order to identify longer-range chromatin interaction loops, we used Mustache, which utilizes a computer vision strategy to identify blob-shaped objects in Hi-C contact matrices ^12^. This approach identified 8,612 loops in the cortex and 8,971 loops in the medulla. Next, we used Mustache to identify intrachromosomal loops with significantly different contact density between the cortex and medulla. This identified 2,535 loops that showed stronger contact density in the cortex and 3,306 loops that preferentially existed in the medulla. The length of the loops in the cortex (mean = 363,629 bp, standard deviation = 313,790 bp, mode = 120,000 bp) and medulla (mean = 379,997 bp, standard deviation = 313,364 bp, mode = 120,000 bp) demonstrated similar, positively skewed distributions (Supplementary Fig. 7a). Consistent with previous reports ^12,13^ a significant proportion of the loop termination points in the cortex (72.14%) and medulla (74.07%) demonstrated overlap with CTCF binding domains (Supplementary Fig. 7b). Consistent with a role for loop termination points in actively maintaining chromatin architecture, a low percentage of the loop termination points in the cortex (22.78%) and medulla (22.35%) demonstrated overlap with H3K9Me3, a heterochromatin signature ^14^ (Supplementary Fig. 7c).

To assess which of these long-range chromatin interactions/loops were connecting a DAR to a DEG, we extended the loop termination points by ±25kb at both ends to create “loop anchors”. We found that the density of DARs adjacent to significantly differential loops was similar at 1.66/Mb in cortex and 1.61/Mb in medulla whereas the transcriptome wide DAR density was only 0.35/Mb. Similarly, the DEG density adjacent to significantly differential loops was 12.81/Mb for cortex and 12.79/Mb for medulla with transcriptome wide DEG density being only 1.20/Mb. These results showed that both DARs and DEGs were highly enriched in the vicinity of cortex- and medulla-specific chromatin loops.

We then asked which of the highly concordant DEGs with DARs also demonstrated a differential loop. Even with these stringent intersecting criteria, we identified 25 genes with concordant multi-omic support at the levels of differential expression, accessibility and chromatin loops between cortex and medulla (Fig. 4e), Supplementary Table 16). This was confirmed by direct examination of the multi-omic landscape at the exemplar cortex gene loci, *MME, SLC7A7* and *KCNJ15*, (Supplementary Fig. 8) and the exemplar medulla gene loci *SLCO3A1, CLDN14* and *WNT7B* (Fig. 5). Many kidney trait related GWAS variants concentrated in accessible chromatin regions close to differentially expressed genes in the medulla (e.g., for *CLDN14 and WNT7B*). In some cases GWAS variants could be linked to a target gene via a medulla-specific chromatin loop (e.g., for *SLCO3A1*). These data confirmed that DARs and chromatin loops were correlated with DEGs and could serve to dissect the cortex and medulla-specific regulatory programs driving gene expression and genetic susceptibility to complex kidney-related traits (see below).

**Figure 5:**
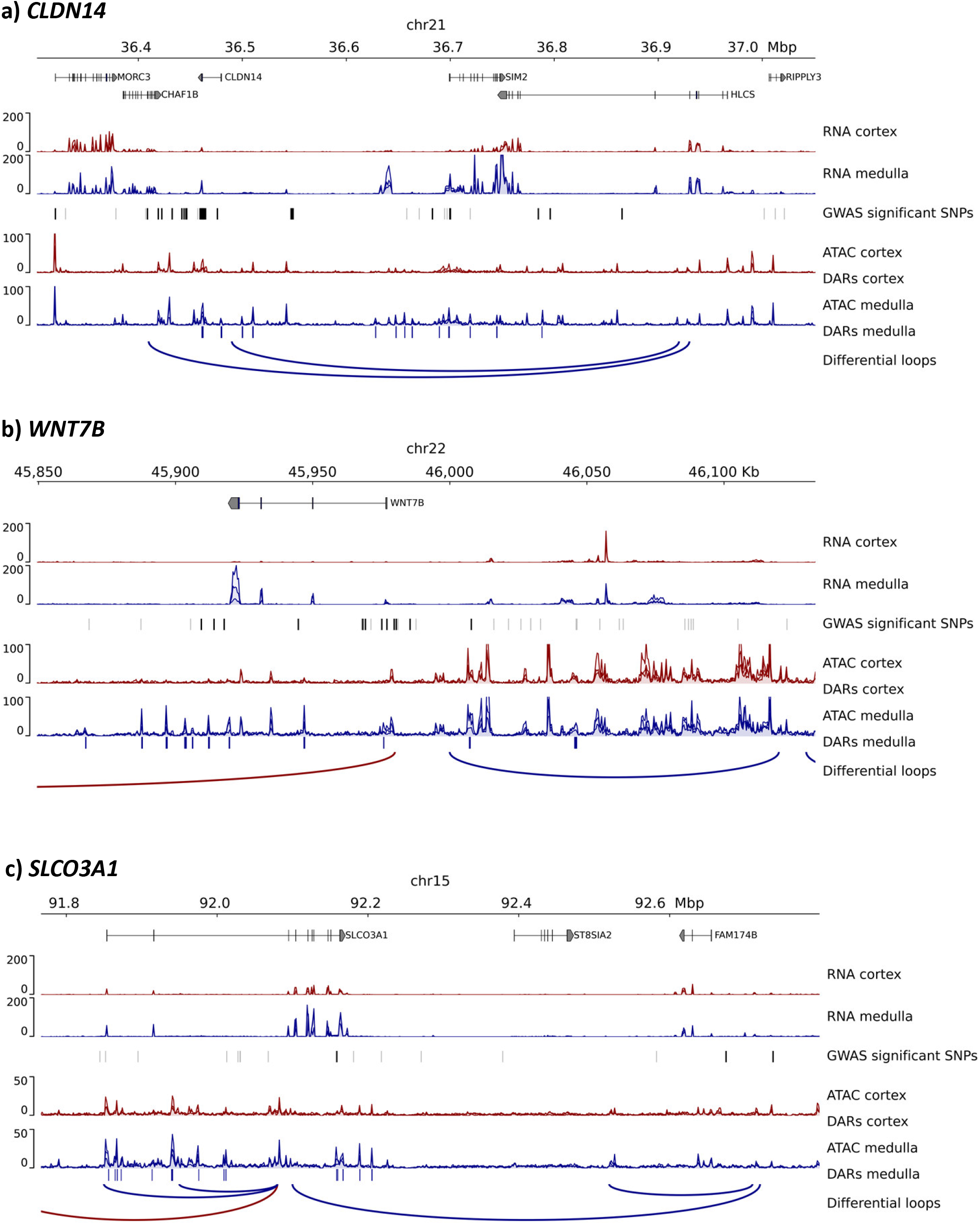
Epigenomic landscape around exemplary medulla genes. Normalized RNA-seq and ATAC-seq signal for our three cortical and medullary samples (overlay of three samples for each track). DARs are marked with boxes (DARs cortex / medulla) below the respective signal tracks. Differential loops from Hi-C data are color coded according to preferential signal in cortex (red) or medulla (blue). The landscape around **a)** *CLDN14*, **b)** *WNT7B* and **c)** *SLCO3A1* is shown. For each region, all significant GWAS SNPs from the NHGRI-EBI GWAS Catalog are displayed (black: SNPs associated with kidney related traits, grey: All other SNPs).

### Identification of transcription factor drivers of the medullary gene expression program

To identify transcription factors (TFs) driving medullary gene expression, we employed a stepwise approach (Methods). We used all 661 transcription factors from the JASPAR core vertebrate’s database ^15^ as a starting point, and found that 519 out of these (78.5%) were at least minimally expressed in kidney tissue (>= 5 transcripts in >= 2 samples in the intersection of our own and the GTEx dataset). A much smaller set of 35 TFs were differentially expressed in medulla and cortex in both our and the GTEx data. TFs such as HNF1A, WT1, LMX1B and others with known roles in cortical gene expression showed the expected higher expression in kidney cortex and conversely, 23 TFs exhibited significantly higher expression in medulla (Fig. 6a and b). Mutations in *TBX18*, the TF with the most prominent medulla-specific expression pattern, have been linked to dysregulated ureter development and urinary tract malformations ^16^. However, the roles of most of the remaining medulla-specific TFs have not been described. We next sought to identify which of these TFs might be playing a role in medullary gene expression by using two widely used motif enrichment tools (Methods). First, we asked if the binding motifs of JASPAR core vertebrate TFs were enriched in medulla over cortex DARs, or vice versa, using HOMER. For cortex, HOMER identified 109 TFs with significant motif enrichment in cortex DARs and 88 TFs in medulla (Supplementary Table 17). Next, we compared TF motif accessibility across all accessible regions between medulla and cortex samples using chromVar ^17^. This identified 39 TFs with significant accessibility deviations for cortex, which again included HNF1A, HNF1B, VDR, with well-known roles in cortex gene regulation. Conversely, chromVar identified 64 TFs with higher motif accessibility in the medulla. Intersecting these distinct approaches for TF motif enrichment identified 29 significantly enriched TFs in the medulla (Fig. 6c). Finally, we stratified these TFs further to identify those with the maximum level of support across multiple datasets and analysis approaches. This stringent intersectional approach was validated by the identification of HNF1A and VDR as the two TFs with the maximum level of multi-omic evidence in cortex datasets (Supplementary Fig. 9). Conversely, this approach identified POU3F3 as the TF with the maximum level of support across multiple medulla multi-omic datasets (Fig. 6a).

**Figure 6:**
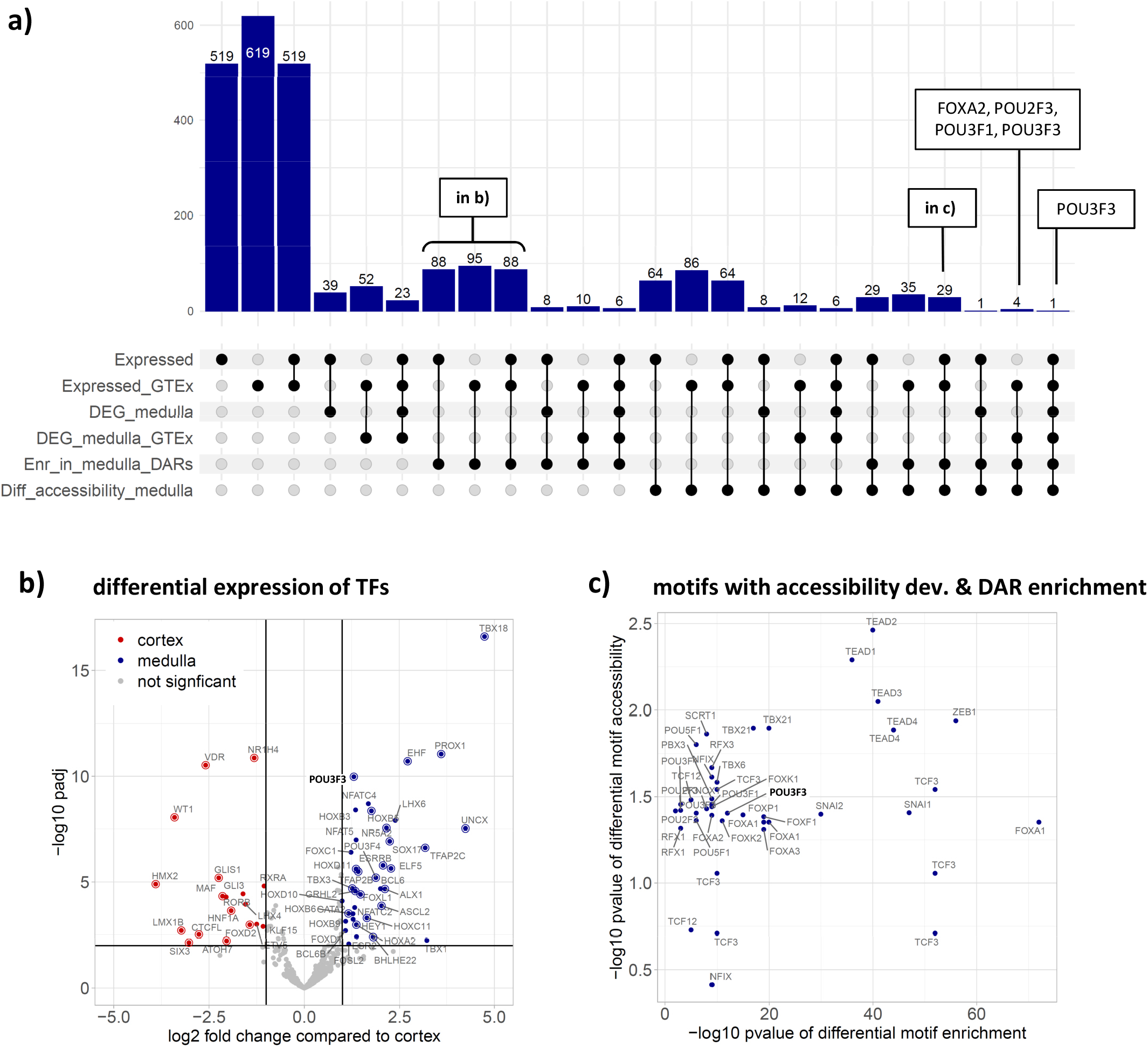
Identification of TFs driving medullary gene expression. **a)** 519 of all 661 TFs in the JASPAR 2022 CORE vertebrates’ database showed at least minimal expression in our and the GTEx V8 kidney samples. The upset plot shows the number of TFs which are furthermore differentially expressed in medulla in our and/or the GTEx data (DEG_medulla, DEG_medulla_GTEx), show significant motif enrichment in medullary DARs (Enr_in_medulla_DARs) and show significant medullary accessibility deviations at their motifs (Diff_accessibility_medulla). Only POU3F3 was identified across all analyses. **b)** Volcano plot showing TFs that are differentially expressed between cortex and medulla in our samples (padj < 0.01, l2fc > 1, indicated by black lines). TFs also showing consistent differential expression in the GTEx data are circled. **c)** Comparison of p-values from differential motif enrichment and accessibility deviations.

### POU3F3 is a key transcription factor in adult and developing kidneys

Our integrative search for medullary TFs converged on POU3F3 (also known as BRN1), which has previously been implicated in fetal kidney development in mouse and human ^18–22^. However, the expression of POU3F3 in adult human kidney medulla has not been studied. We sought to identify putative POU3F3 target genes by first examining whether POU3F3 motifs were enriched in accessible chromatin regions within the promoter region (−5000/+1000 bp from TSS) of DEGs in our own data. Indeed, POU3F3 was the most highly enriched motif in accessible chromatin within the promoter regions of DEGs (Fig. 7a, left). Other POU family members were also enriched within these regions, likely due to similarity of their binding motifs, and this was also the case when looking at motif enrichment in all ARs (Fig. 7a, middle). However, when we examined the gene expression of the other POU-family TFs with similar binding motifs, only POU3F3 was found to be differentially expressed between cortex and medulla (Fig. 7a, right). This made POU3F3 the most plausible family member responsible for POU-motif enrichment in the promoters of DEGs. A similar analysis performed on cortex DEGs promoters identified a key role for HNF1A, as expected (Supplementary Figure 10). Next, we confirmed that POU3F3 was expressed at the protein level in human kidney tissues using immunohistochemistry (Fig. 7c, Supplementary Fig. 11). In the cortex, only distal tubules and collecting ducts expressed POU3F3 within their cytoplasm, with stronger staining of the nucleus; cells in the glomeruli and proximal tubules did not express POU3F3. In the medulla, all tubular components expressed POU3F3 including thick and thin loops of Henle and collecting ducts. Within the medulla, cells in the interstitium and endothelial cells lining medullary capillaries did not express POU3F3.

**Figure 7:**
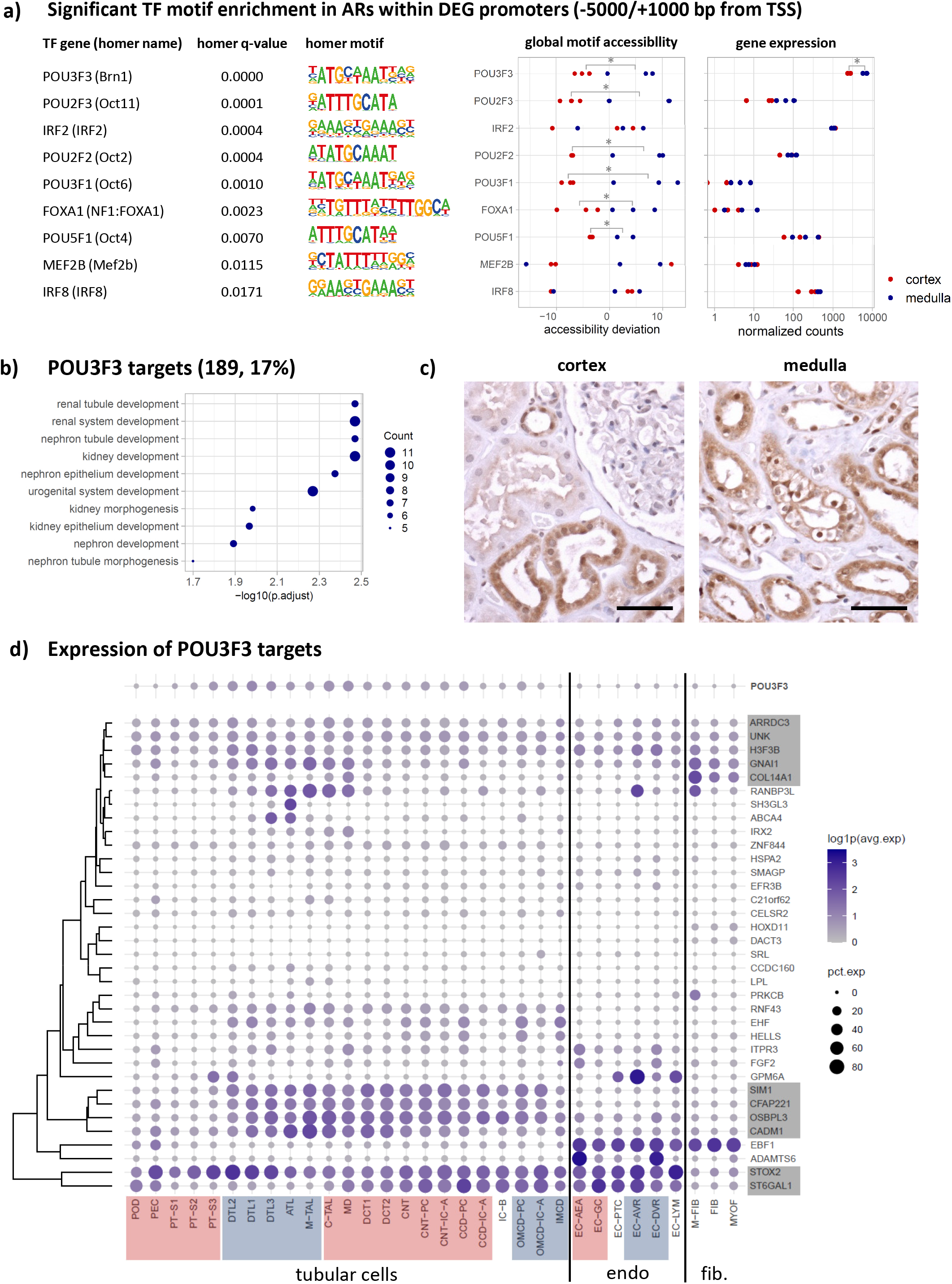
POU3F3 and potential medullary targets. **a)** TFs with significant motif enrichment in ARs inside of medullary DEG promoters (−5000 bp – 1000 bp from TSS) computed using HOMER. The middle panel shows the accessibility deviations for these same TFs computed with chromVar for each of our cortical and medullary tissue samples. The right panel displays the TFs’ normalized expression for each sample. Only POU3F3 is significantly and differentially expressed. **b)** GO-term enrichment of putative POU3F3 downstream target genes. **c)** Immunohistochemical staining for POU3F3 in adult human kidney cortex and medulla. Scale bar, 50µm. **d)** Expression of POU3F3 and its putative targets in the KPMP snRNA-seq dataset. Tubular cell types are ordered along the nephron (red shading: cortical cell types, blue shading: medullary cell types, white if not location-specific). The downstream genes were clustered hierarchically by average expression (grey shading: genes with concordant expression patterns to POU3F3). Only protein coding genes expressed in at least 10% of the cells in at least one cell type were considered for this analysis.

Having confirmed the medulla-enriched expression of POU3F3, we examined the function of medulla DEGs with at least one accessible POU3F3 motif in their promoter (Supplementary Table 18) and found that this set of target genes was enriched for molecular functions associated to kidney development (Fig. 7b). Finally, we sought to correlate *POU3F3* and its target genes’ expression using the KPMP kidney single nucleus gene expression atlas. Confirming our immunohistochemistry staining profile, *POU3F3* was preferentially expressed in tubular components of the distal nephron and medulla in the KPMP atlas (Fig. 7d). Next, we selected the putative POU3F3 target genes with at least 10% expression in any tubular, endothelial or fibroblast cell type and clustered them by average expression per cell type. While some putative target genes were expressed in multiple cell types, a subset (grey shaded boxes) showed expression patterns that were concordant with *POU3F3*. For example, the transcription factor gene *SIM1* has a similar pattern of expression as *POU3F3* in the renal collecting system of the developing mouse kidney ^23^ and pronephros of zebrafish ^24^. Another potential POU3F3 target gene, *RANBP3L* has been shown to play a role in BMP signaling pathways ^25^, ischemia reperfusion injury ^26^ and has been linked to regulation of blood pressure via urinary potassium excretion ^27^. *CADM1*, which has a distal nephron/medullary pattern of expression overlapping with *POU3F3* ^28^, appears to be shed in nephropathies ^29^ and may serve as a urinary biomarker of medullary injury ^30^. Therefore, our analysis of genes with POU3F3 motifs within their promoter identified several genes with plausible roles in kidney development and disease.

## Discussion

We have previously generated chromatin accessibility (ATAC-seq) and conformation data (Hi-C) to identify the regulatory control mechanisms of cultured kidney cells ^31^. While useful, *in vitro* culture can induce injury responses in cells ^32^, and therefore characterization of intact tissues with minimal manipulation is essential to understand their baseline epigenomic landscapes. Here we present reference-quality functional genomic datasets for human kidney cortex and medulla that will provide a valuable resource for kidney investigators. With careful histologic curation and evaluation of bulk gene expression signatures, we were able to assign GTEx kidney samples to cortex and medulla. This identified some misassigned samples in this publicly available resource, which had prevented meaningful differential gene expression analysis. However, after sample reassignment, we were able to extract medulla-specific gene expression signatures from the GTEx kidney samples that were not previously appreciated and contained numerous positive controls. Our approach therefore may have implications for other tissues available through GTEx, and could be applied to those that exhibit evidence of sample misassignment ^33^.

Integrative analyses identified several principles of gene regulation in the kidney cortex and medulla: differentially accessible chromatin regions in promoters were positively correlated with differential expression of the nearest gene. Structural differences in chromatin architecture (i.e., loops) between cortex and medulla were also correlated with differentially accessible chromatin regions and differentially expressed genes. Integrating all levels of information across our own and the GTEx samples, we identified a set of 71 genes characterizing the distinct epigenomic landscapes of the human kidney cortex and medulla. Metabolic and transporter functions were predictably enriched in genes preferentially expressed in the cortex. By contrast, genes related to angiogenesis and extracellular matrix were enriched in the medulla. The implication of genes related to angiogenesis may reflect the hypoxic milieu of the renal medulla, whereas identification of extracellular matrix genes is consistent with the medulla’s distinct histologic appearance and composition during development ^34^. Even so, the distinct matrix composition of the medulla has not been systematically studied in the adult.

The exemplar medullary genes also revealed the need for more detailed investigation of the medulla and its contribution to kidney disease. For example, the medullary gene *CLDN14* encodes a tight junction protein involved in paracellular calcium reabsorption ^35^. *CLDN14* is highly expressed in cells of the medullary thick ascending limb of the loop of Henle in single-nucleus RNA-seq data from human kidney ^7^. Common genetic variants in *CLDN14* have been associated with kidney stone disease ^36^. Other exemplar genes are poorly characterized in the kidney. For instance, *SLCO3A1* encodes an organic anion transporter expressed in multiple human tissues with high and selective expression in the medulla. Our data demonstrate numerous medulla-specific accessible chromatin regions associated with medullary chromatin loops across the *SLCO3A1* promoter and gene body. SLCO3A1 has been implicated in Crohn’s disease and experimental colitis ^37,38^, cholestasis ^39,40^, and Parkinson’s disease ^41^. Therefore, determining its role in the adult kidney medulla will be an intriguing future line of investigation.

Interestingly, the gene ontology analyses also highlighted genes associated with kidney development as enriched in the medulla. This suggests that these genes also play an important role in the adult medulla, beyond a role in development. Consistent with this idea, our integrative analysis identified *WNT7B*, a gene crucial for medulla development during embryonic stages ^42^ but also described to enhance renal tubule repair after ischemic injury ^43^. Our investigation of transcription factor binding motif enrichment in medulla open chromatin regions identified POU3F3, a highly conserved transcription factor ^19^ that plays a critical role in development of the excretory system in worms ^44^, as well as in the distal nephron of frogs ^45^ and mice ^18–20^. Single cell gene expression studies showed the highest expression of *POU3F3* is in the developing loop of Henle in mice ^21^ and humans ^22^. In the adult human kidney, *POU3F3* is expressed in the loop of Henle and distal tubule, but also in the cortical and medullary collecting ducts ^7^, a localization pattern that differs slightly from that in mice. We were able to confirm POU3F3 protein expression at these sites in adult human kidney tissues by immunohistochemistry. POU3F3 target genes are enriched in kidney and urogenital development gene ontologies, and our results implicate a role for them in the functioning of the adult kidney medulla. Based on our findings, the role of POU3F3 expression in the adult human kidney medulla and its contribution to kidney disease will be an exciting area for future investigation.

In summary, we show that generation of carefully curated reference data sets allow for reassignment of tissue labels in publicly available RNA-seq data, which may also have implications for GTEx tissues outside the kidney. Our epigenomic maps of the adult human kidney reveal a distinct genome organization, chromatin accessibility landscape, and gene expression profile for the renal medulla as compared to cortex. These maps will serve as a valuable reference for studies on kidney gene regulation, genetic susceptibility to disease including GWAS, and the contributions of the medulla to normal kidney function and disease.

## Methods

### Tissue dissection and RNA sequencing

Kidney samples were obtained from macroscopically dissected cortex and medulla of tumor-adjacent normal tissue in nephrectomy specimens from three donors as described previously ^46^. Briefly, RNA extraction and RNA-seq was performed by GeneWiz. Trimming and alignment of paired-end fastq files to human reference genome sequence hg38 was done with STAR 2.7.5b ^47^ with parameters --outFilterIntronMotifs RemoveNoncanonical --outFilterMismatchNoverReadLmax 0.04. Counting of the number of reads aligned to each exon (feature) was performed using *featureCounts* ^48^.

### Tissue decomposition with BisqueRNA

In order to estimate the cell-type compositions in the cortex and medulla samples, we applied the tool Bisque ^8^ to decompose our bulk RNA-seq data using snRNA-seq data from KPMP as reference. The snRNA-seq data was retrieved from the KPMP Kidney Tissue Atlas at https://atlas.kpmp.org/explorer/dataviz ^7^ and processed by Seurat package (v4.1.0) ^49^. Genes were clustered based on average expression (Euclidean distance). Within the R package BisqueRNA (version 1.0.5), we used the function ReferenceBasedDecomposition with default parameters for decomposition.

### Differential gene expression analysis of our samples

Differentially expressed genes were computed with DESeq2 ^9^ with default parameters. Only transcripts with a count of at least 5 in >= two samples were selected for the analysis. Shrunken log2 fold changes were generated with the lfcShrink function (“apeglm” method) to make log2 fold changes comparable across a wide range of counts. All genes with a log2 fold change > 1 and an adjusted p-value < 0.01 were considered differentially expressed. Absolute count differences for each transcript were derived from the respective base mean and log2 fold changes. GO term over-representation analyses were performed using the enrichGO function from clusterProfiler (3.18.1) ^50^ with all transcripts included in the DESeq analysis as background. GO term enrichment analyses for the GTEx data were carried out identically.

### Reassignment and differential gene expression analysis of GTEx V8 samples

The gene read counts data (GTEx_Analysis_2017-06-05_v8_RNASeQCv1.1.9_gene_reads.gct.gz) and sample attributes (GTEx_Analysis_v8_Annotations_SampleAttributesDD.xlsx, including pathology notes) were retrieved from https://www.gtexportal.org/home/datasets, and the RNA-seq data from kidney samples was extracted by filtering for SMTS == “Kidney” & SMAFRZE == “RNASEQ”. Transcripts with a count of at least 5 in >= 2 samples were selected, then the counts were variance stabilized and normalized for library size using the DESeq2 vst function. The PCA was computed using the DESeq2 plotPCA function with default parameters. For the combined PCA with our own samples, we first selected transcripts with a raw count of at least 5 in >= 2 of our own samples and then performed an inner join with the GTEx data based on Ensembl IDs in order to select transcripts which were detected in both datasets. The combined count matrix was variance stabilized and normalized for library size using the DESeq2 vst function. The combined PCA was computed with the DESeq2 plotPCA function using the top 1000 most variable features. GTEx samples were reassigned based upon proximity to our own medulla and cortex reference samples in the combined PCA. Specifically, all GTEx samples that 1) mapped to the left of our own right-most sample scored as 100% medulla by a trained pathologist on PC1, and 2) were located on the same side of 0 as all of our own medullary samples on PC2 were assigned as “medulla”. The remaining GTEx samples were assigned “cortex”. The differential gene expression analysis with GTEx samples before and after reassignment was carried out identically to the one of our own samples.

### ATAC sequencing and bioinformatics workflow

ATAC-seq was carried out on snap-frozen human kidney samples by ActiveMotif (Carlsbad, CA) and aligned to the hg38 reference genome (BWA default settings). Only properly aligned read pairs with a mapping quality >= 40 aligning to the same chromosome were selected with samtools 1.13 ^51^. Read duplicates were removed with picard 2.25.5, and reads located within the ENCODE blacklist regions were removed with deepTools alignmentsieve 3.5.1 ^52^. Accessible regions (ARs, peaks) for each sample were called with MACS 2.1.0 ^53^ at a cutoff q-value of 0.01 without control file and the –nomodel, -shift −100 and –extsize 200 options. A masterlist of ARs across all samples was created using bedops –m (bedops 2.4.35 ^54^). Then normalization of read depth was carried out by random down-sampling to the sample with lowest coverage. The counts for all master list ARs for all samples were determined with bedops bedmap –count. Differentially accessible regions (DARs) and log2 fold changes (l2fc) in counts for accessible regions between cortex and medulla were identified with DESeq2 (default settings). All regions with an adjusted p-value < 0.05 and a l2fc > 0 were considered to be differentially accessible. PCA plots with all samples were generated from the Rlog transformed count data for each AR in the masterlist with the DESeq2 plotPCA function. ARs and DARs were assigned to genes and the respective genomic features based on proximity with the ChIPseeker 1.28.3 annotatePeak function ^55^. Promoters were defined as the region −5000 to 1000 bp from the TSS, downstream was defined as the region less than 5000 bp downstream of the gene body. The genomicAnnotationPriority parameter was set to the following order: “Promoter”, “Exon”, “Intron”, “5UTR”, “3UTR”, “Downstream”, “Intergenic”. The ChIPseeker based assignment of ARs and DARs to genes was used to test for enrichment of DARs within DEG promoters (Fisher’s exact test, fisher.test function from R 4.1.0 stats package)(ChIPseeker reference: PMID: 25765347). The Pearson correlation coefficients for log_2_-fold changes in AR counts and gene expression were computed with the cor.test function from the R stats package.

### Identification of DEGs with DAR enrichment

Genes with enrichment of DARs (FDR < 0.05) were identified with GREAT 4.0.4 ^56^ (Species assembly: hg38; Association rule: Basal+extension: 5000 bp upstream, 1000 bp downstream, 0 bp max extension, curated regulatory domains included) using either the cortical or medullary DARs as test region and the non-differential ARs as background. DEGs with DAR enrichment for cortex and medulla were the identified by overlapping the respective lists of DEGs and genes with DAR enrichment for cortex and medulla. The subset of DARs with DEG enrichment in proximity of differential Hi-C loops was identified by extending the Hi-C loop anchors by 25 kb in both directions and overlapping them with the respective gene bodies (overlapsAny function, GenomicRanges version 1.44.0 ^57^).

### Regional plots for selected genes

Visualization of read-normalized density tracks (.bw) from the RNA-seq and ATAC-seq experiments along with DAR regions (.bed) and differential loops (.bedpe) was done with pyGenomeTracks version 3.7. ^58^ using the NCBI RefSeq select genome annotation. GWAS variants for the displayed genomic intervals were downloaded from the NHGRI-EBI GWAS Catalog^59^ website (https://www.ebi.ac.uk/gwas/) and filtered for genome wide significance (p-value <= 5×10e-8). All significant variants were categorized manually into kidney-related and non-kidney-related (other) based on their associated trait. Kidney-traits included all kidney function parameters (e.g. eGFR, UACR), kidney disease traits (e.g. kidney stones) electrolyte levels, traits associated with electrolytes (e.g. bone mineral density), levels of drugs and metabolites with renal elimination, traits associated with fluid balance, blood pressure and blood formation (e.g. systolic blood pressure or hematocrit) and traits associated with glucose metabolism (e.g. type 2 diabetes).

### TF motif enrichment in DARs

Transcription factors (TFs) with motif enrichment in cortical and medullary DARs were identified with HOMER v4.11 findMotifsGenome (options: –size given –mask) ^60^. Known TFs with a q-value below 0.05 were considered significant. For motif enrichment in cortical DARs, the medullary DARs were used as background and vice versa. Both sets of regions had nearly identical size distributions based on mean, median and histogram, hence the option “–size given” could be chosen for findMotifsGenome. The HOMER names of TFs with significant enrichment were mapped manually to the names used in the JASPAR CORE database for comparability with the chromVar. This mapping is shown in Supplementary Table 14.

### TF motif enrichment in accessible promoter regions

The TSS for all DEGs based on their canonical transcript was retrieved from Ensembl (genomic build GRCh38.p13) via biomaRt 2.48.3 ^61^. Promoters were defined as the region – 5000 bp to 1000 bp from the TSS. The merged set of ARs identified in the three cortical and the three medullary samples was screened for any ARs overlapping the promoters of cortical or medullary DEGs. HOMER v4.11 findMotifsGenome (options: –size given –mask) was used to identify TFs with motif enrichment in ARs of medullary and cortical DEG promoters. For cortex, the ARs in medullary DEG promoters were used as background, and vice versa. TFs with a q-value below 0.05 were considered significant.

### TFs with differential motif accessibility

TFs with differential motif accessibility were identified with chromVar 1.14.0 ^17^. The analysis was based on the merged set of ARs identified in the three cortical and the three medullary samples. All ARs were resized to an equal width of 500 bp around the AR center. Read depth for all samples was randomly down-sampled to match the sample with lowest coverage. chromVar getCounts was used to obtain the fragment counts per accessible region. Transcription factor motifs were retrieved with the JASPAR2020 R-package ^15^ and limited to the JASPAR CORE vertebrates motif set. ARs containing the respective TF motifs were annotated with motifmatchR 1.14.0 and then used for computing the accessibility deviation between cortex and medulla for each TF with chromVar. TFs with a deviation p-value < 0.05 were considered significant for either cortex or medulla, depending on if the mean deviation was higher for either for cortex or medulla.

### Combined approach to identify medulla and cortex specific TFs

All TFs from the JASPAR CORE vertebrates database were marked for expression (count > 5 in >= 2 samples), differential expression (p-adj 0.01, l2fc > = 1), significant motif enrichment in DARs (q-value < 0.05) and differential accessibility (p-val < 0.05) in either cortex or medulla. The number of TFs meeting different subsets of criteria were computed and depicted with ComplexUpset 1.3.3 ^62^.

### Identification of putative POU3F3 downstream target genes

All medullary DEGs were screened for POU3F3 motif occurrences within ARs located in their promoters (−5000 bp to 1000 bp from the canonical TSS) with motifmatchR 1.14.0 using the TF motif MA0788.1 from the JASPAR 2020 CORE vertebrate database (p.cutoff = 5e-5). The same screen was additionally performed with HOMER v4.11 annotatePeaks using the Brn1 motif provided by HOMER. The putative POU3F3 downstream targets from both screens were then filtered for protein coding genes.

The expression in selected kidney cell types was only shown for genes identified in both motif searching approaches and for genes expressed in at least 10% of the cells in any of the cell types. The snRNA-seq data was retrieved from the KPMP Kidney Tissue Atlas at https://atlas.kpmp.org/explorer/dataviz ^7^. Genes were clustered based on average expression (Euclidean distance).

### Chromosome conformation data (Hi-C)

Hi-C was performed on macro-dissected kidney cortex and medulla from a 74-year-old man. Approximately 0.5 cm^3^ of tissue was minced with a razor blade and fixed and nutated for 20 minutes in 10 mls of a 1:10 dilution of 10% neutral buffered formalin (Fisher). Following this, the formalin was quenched by direct addition of 0.1 g of glycine (Sigma) and nutation for another 15 minutes. The tissue fragments were submitted to Phase Genomics (Seattle, WA) for Hi-C library preparation using their Human Hi-C Kit and sequencing on an Illumina HiSeq 4000. Following alignment and downsampling to equal read numbers, contact matrices (.hic files) were generated and visualized using the Juicebox application (https://www.aidenlab.org/juicebox/). Topologically associating domain (TAD) calling was performed using the DomainCaller algorithm at 50 kb resolution with default parameters.

Intrachromosomal interactions (loops) in cortex and medulla were inferred using Mustache ^12^ from the .hic files. We used a p-value threshold of 0.1 with a sparsity threshold of 0.7 for detecting intrachromosomal interactions, and an fdr threshold of 0.2 to identify differential intrachromosomal interactions between cortex and medulla. The total numbers of DARs and DEGs in the genomic windows of loop anchors + 25 kb flanking regions were calculated. The results were normalized to density of DARs and DEGs per Mb genomic region. The density of DARs and DEGs adjacent to differential loops were compared against the average density of DARs and DEGs in the total human transcriptome. The bedfiles of CTCF (ENCSR871TQJ:ENCBS136AAA, ENCBS137AAA), H3K9Me3 (ENCSR000HPR, ENCSR378HAM), and H3K27Ac (ENCSR230BWN, ENCSR438SPO) ChIP-seq were downloaded from the ENCODE project website. The proportion of loop anchors overlapping CTCF, H3K9Me3, and H3K27Ac were calculated by counting the number of loop anchors +25 kb flanking regions with at least one overlapping ChIP-seq peak of interest.

### POU3F3 immunohistochemistry

5 µm thick sections of formalin fixed paraffin embedded kidney cortex and medulla were affixed to charged microscope slides and baked at 60°C for 30 minutes. After deparaffinization and rehydration, sections were subjected to heat induced antigen retrieval using citrate buffer, pH 6 for 10 minutes at 110°C in a BioCare Medical decloaking chamber. After blocking, polyclonal rabbit primary antibodies reactive against POU3F3 (Atlas Antibodies, HPA067151) were applied to the tissue sections at 0.5 µg/ml overnight at 4°C. After washing, secondary antibody labeling and chromogenic development were performed using an ImmPRESS horseradish peroxidase Horse Anti-Rabbit IgG PLUS Polymer Kit (Vector Biolabs, MP-7800-15). After development, slides were coverslipped with EcoMount (BioCare Medical, EM897L).

## Supplementary Figures

**Supplementary Figure 1:**
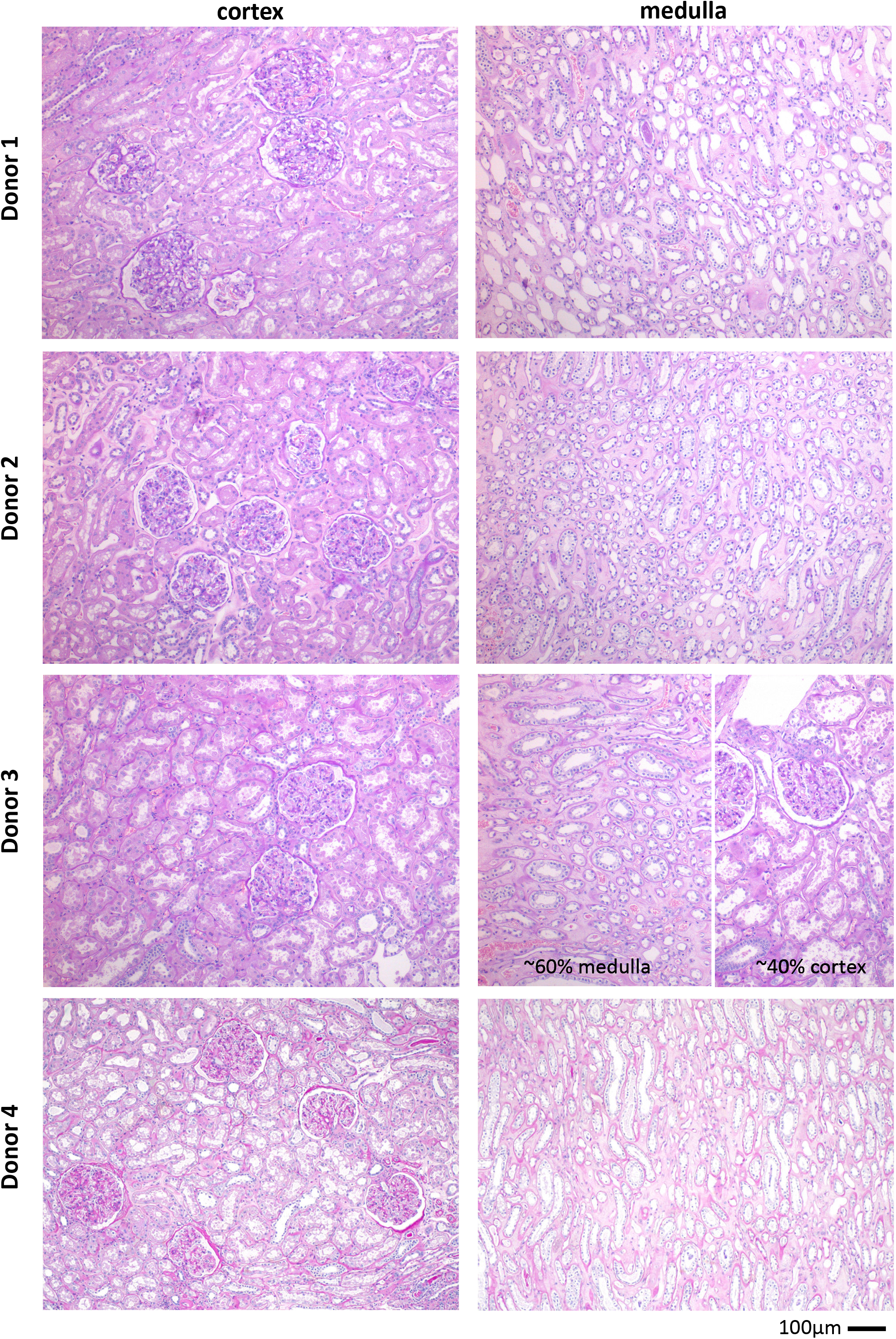
Cortex and medulla samples used for RNA-seq, ATAC-seq and Hi-C (PAS staining)

**Supplementary Figure 2:**
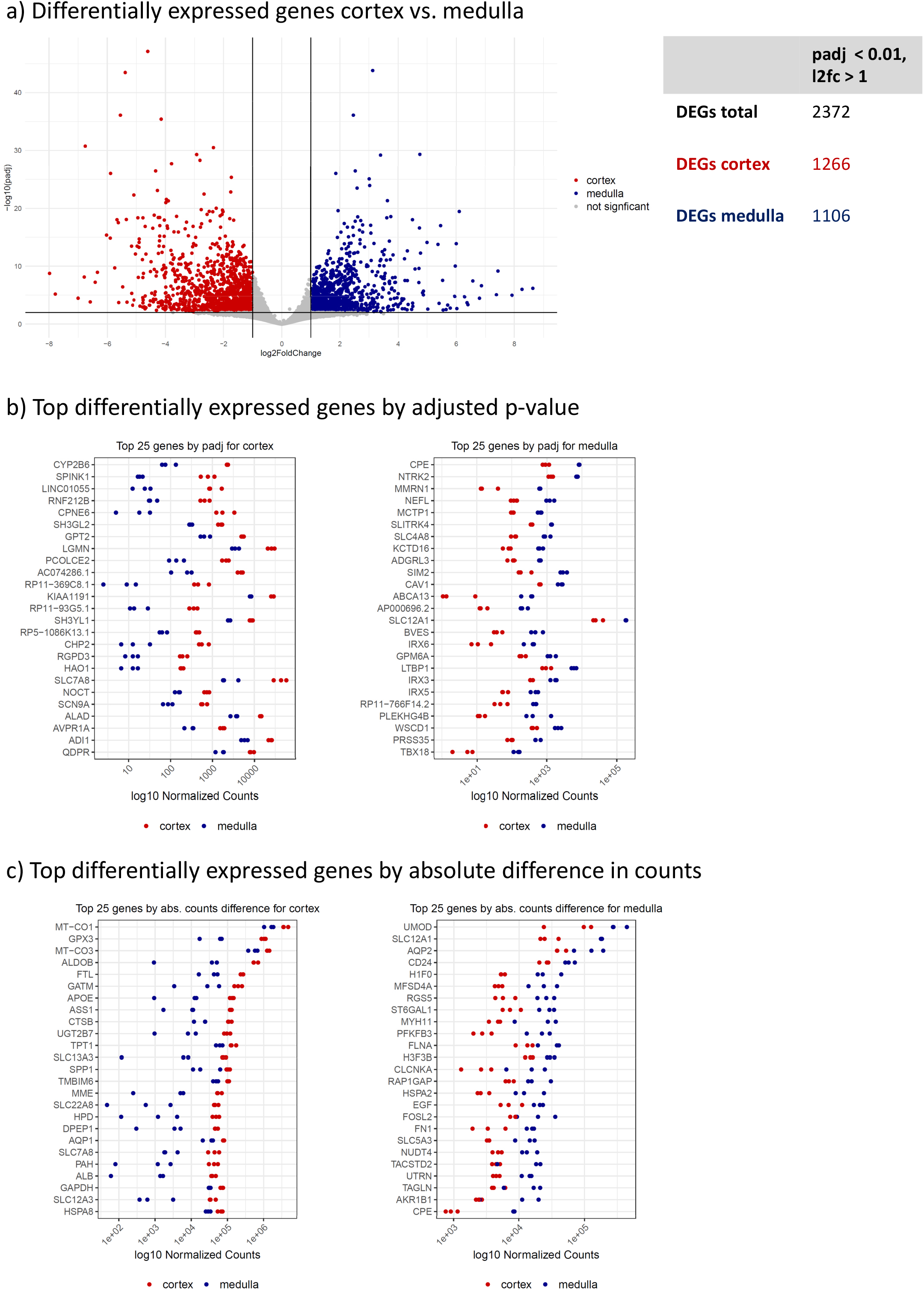
Results from differential gene expression analysis cortex vs. medulla.

**Supplementary Figure 3:**
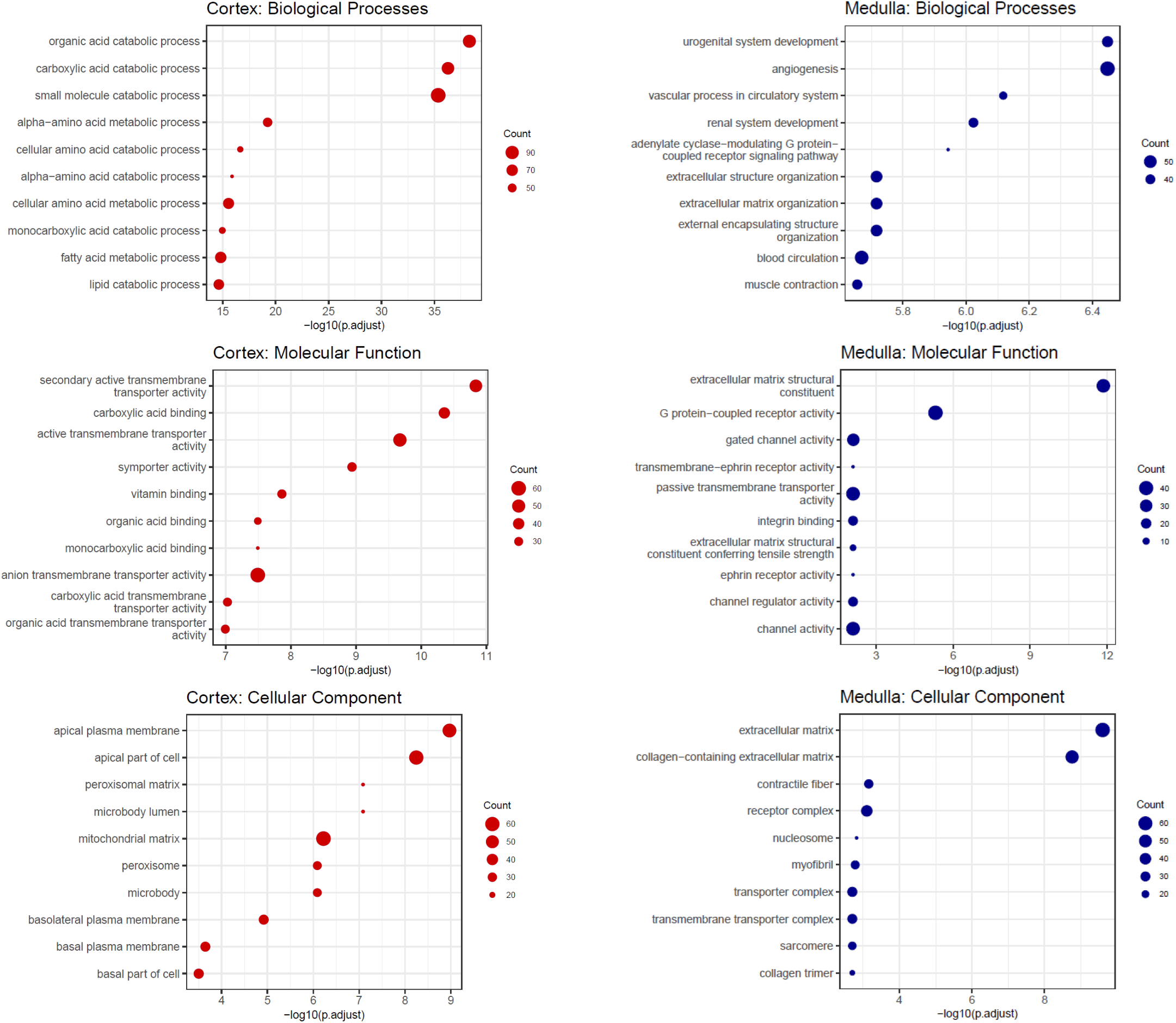
Enriched GO-terms for cortex and medulla DEGs (own samples)

**Supplementary Figure 4:**
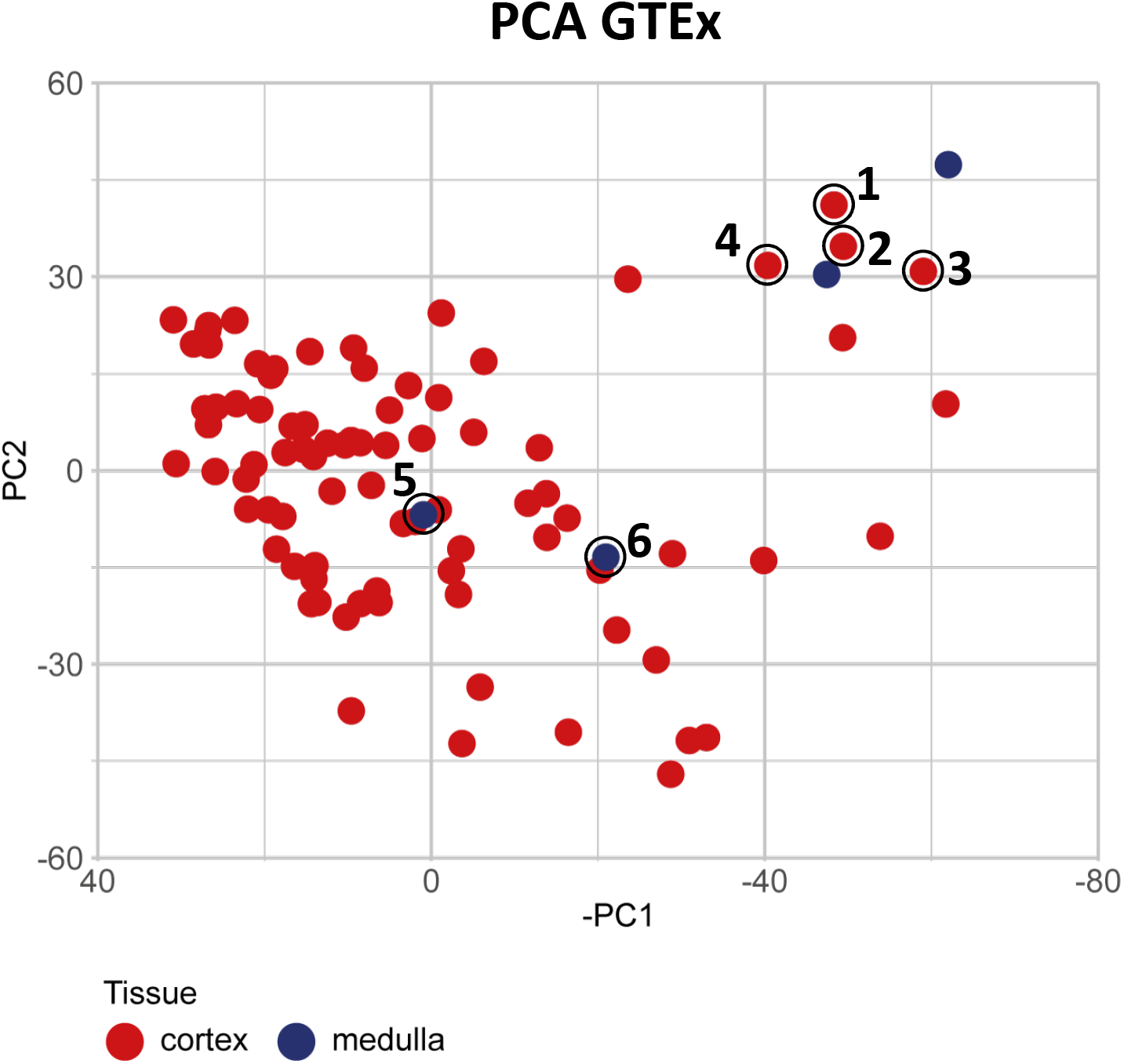
Conflicting GTEx pathology notes. 1) “6 pieces; 2 pieces are mostly medulla” 2) “6 pieces; predominantly cortex, 2 are (largely) medulla, arteriosclerosis” 3) “2 aliquots, ∼8.5×8mm. All cortex, may be used when re-labled as cortex. Tubules autolyzed, crystalloid deposits in tubular lumina” → originally medulla, then relabeled as cortex. 4) “6 pieces, glomeruli present in 5/6; one is mainly medullary rays (delineated).” 5) „2 aliquots, ∼8×6mm. Minor (∼20-30%) ‘contamination’ by cortex with glomeruli, noted on images” 6) “2 pieces, ∼8×5mm. NOTE: Each aliquot is ∼40% cortex”

**Supplementary Figure 5:**
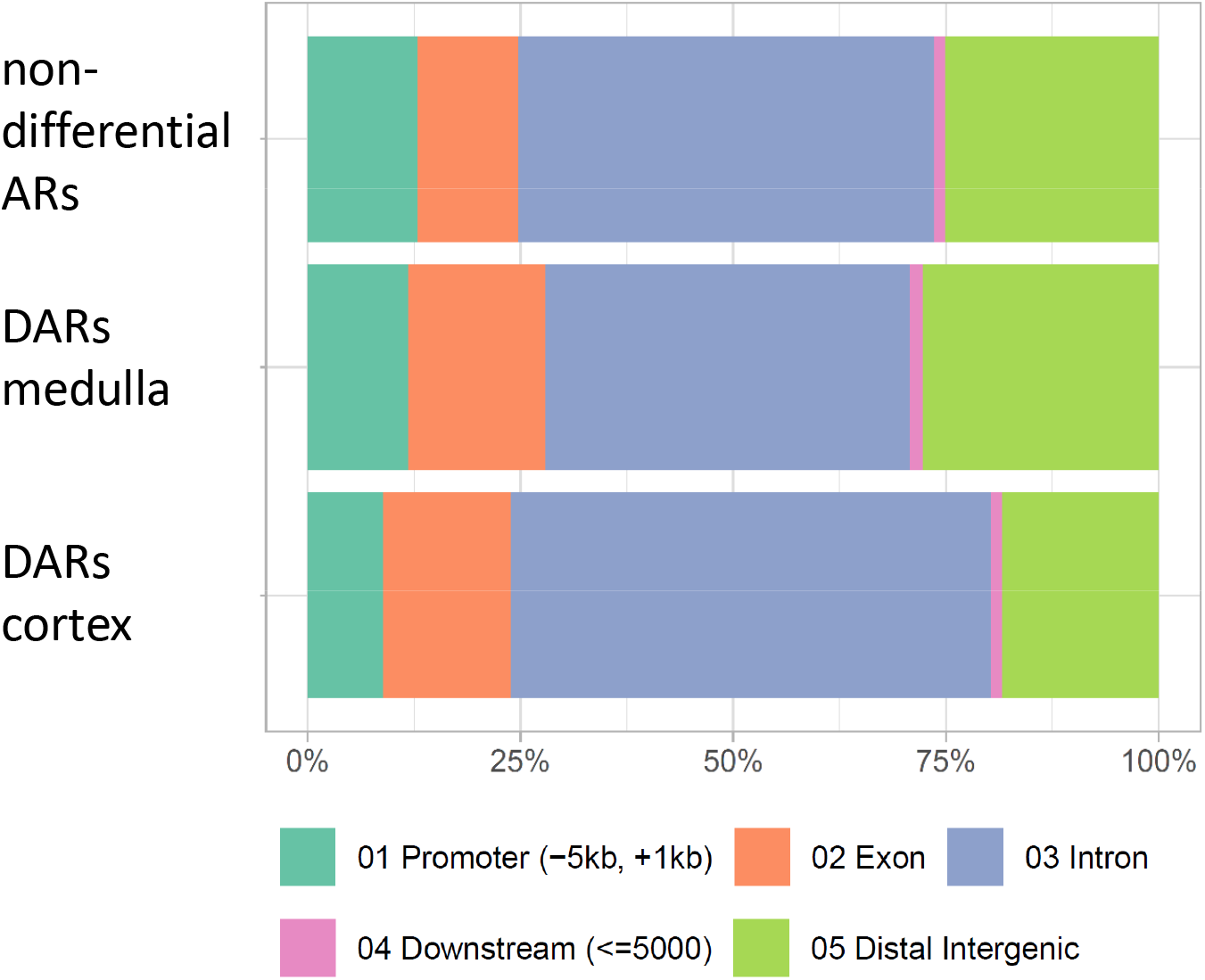
Genomic location of ARs and DARs. Percentage of non-differential ARs and DARs for medulla and cortex located in the different genomic regions (Promoter: −5000 bp to 1000 bp from TSS, Downstream: Transcription end site to 5000 bp downstream, other regions as indicated)

**Supplementary Figure 6:**
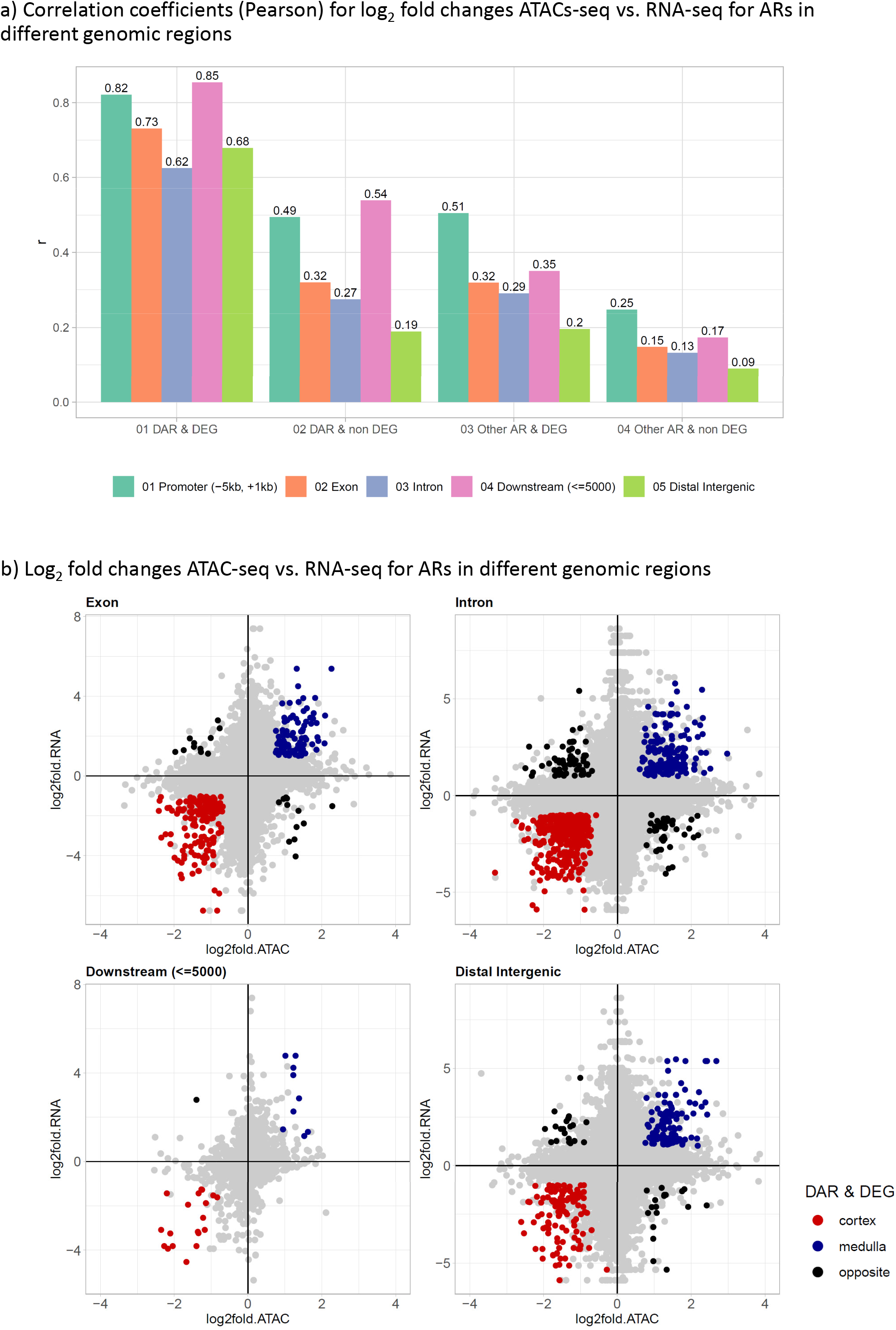
Correlation between log_2_ fold changes of ATAC-seq signal and gene expression.

**Supplementary Figure 7:**
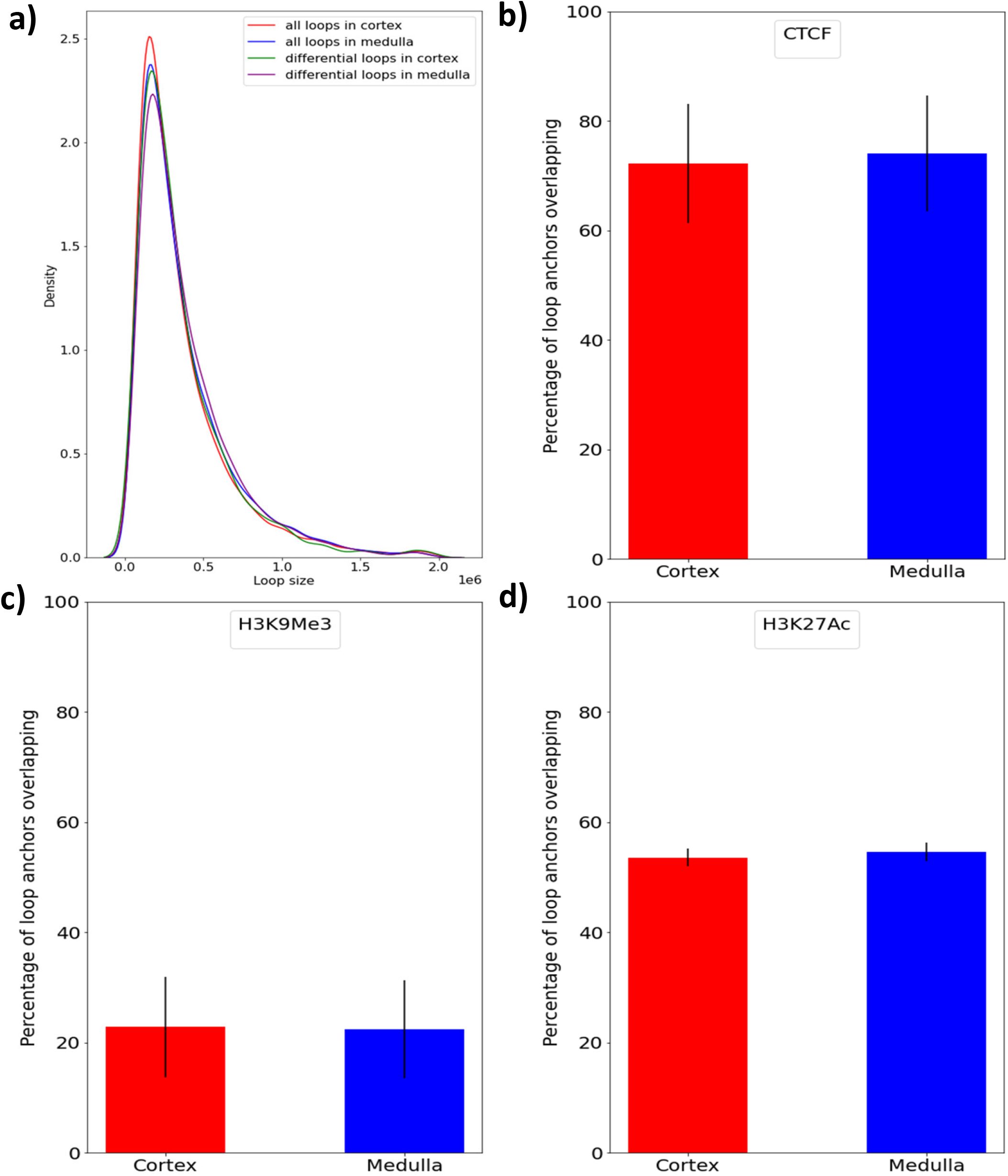
Hi-C loop size distribution and loop overlap with CTCF, H3K9Me3 and H3K27Ac binding sites. Hi-C sequencing demonstrates similar distribution of intrachromosomal interactions in the cortex and medulla (a). A high proportion of the anchors of the intrachromosomal interactions in the cortex (72.14%) and medulla (74.07%) overlap with CTCF binding domains in kidney tissue (b). Interestingly, very low percentage of loop anchors in the cortex (22.78%) and medulla (22.35%) demonstrate overlap with H3K9Me3 (c). Both cortex (53.55%) and medulla(54.61%) demonstrate similar proportion of anchors overlapping H3K27Ac binding regions in the cortex and medulla (d). The error bars indicate standard errors of mean.

**Supplementary Figure 8:**
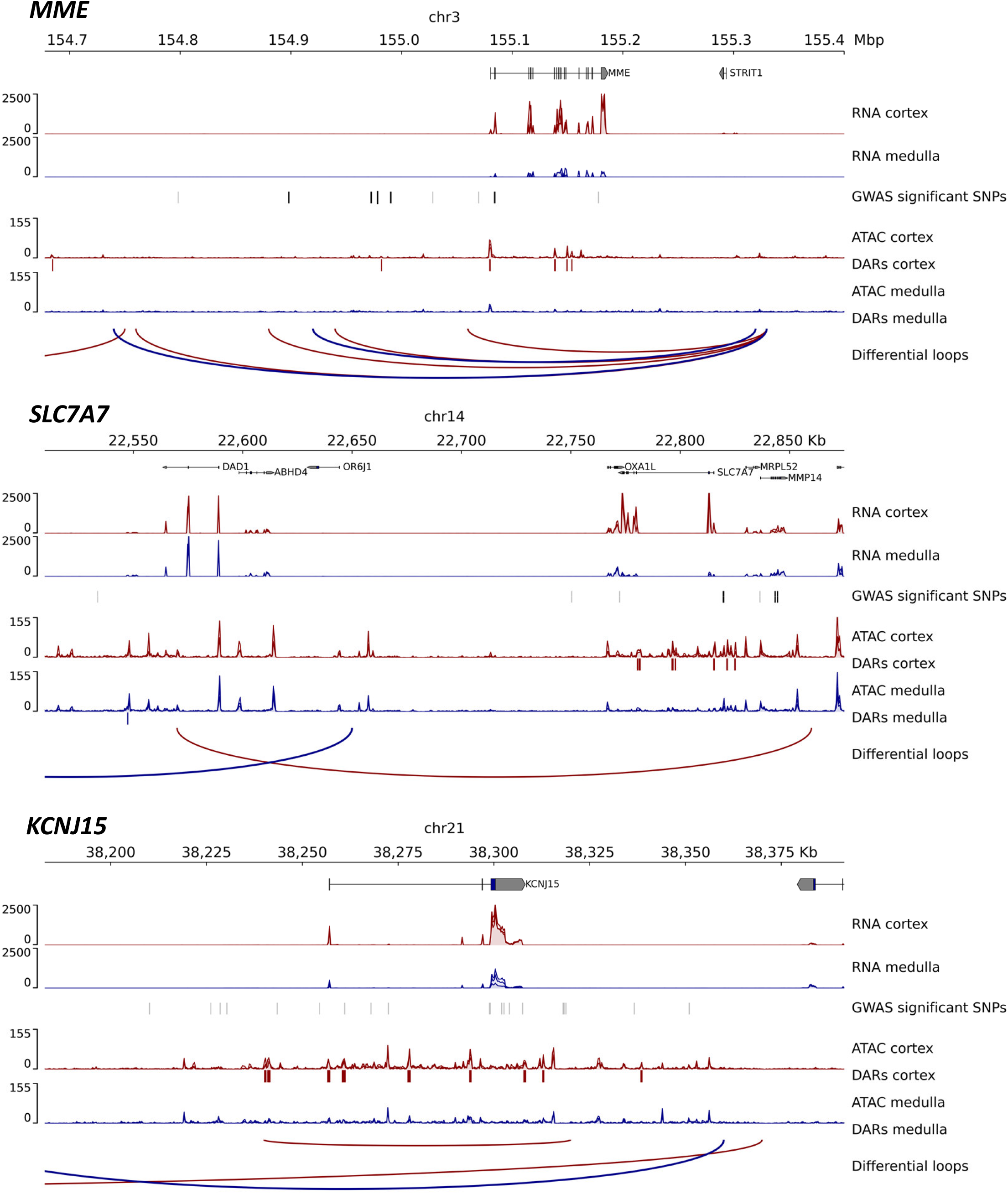
Genomic regions around exemplary genes with higher medullary expression and chromatin accessibility as well as long range chromatin interactions specific for medulla. Normalized RNA-seq and ATAC-seq signal for our three cortical and medullary samples (overlay of three samples for each track). DARs are marked with boxes (DARs cortex / medulla) below the respective signal tracks. Differential loops from Hi-C data are color coded according to preferential signal in cortex (red) or medulla (blue). For each region, all significant GWAS SNPs from the NHGRI-EBI GWAS Catalog are displayed (black: SNPs associated with kidney related traits, grey: All other SNPs).

**Supplementary Figure 9:**
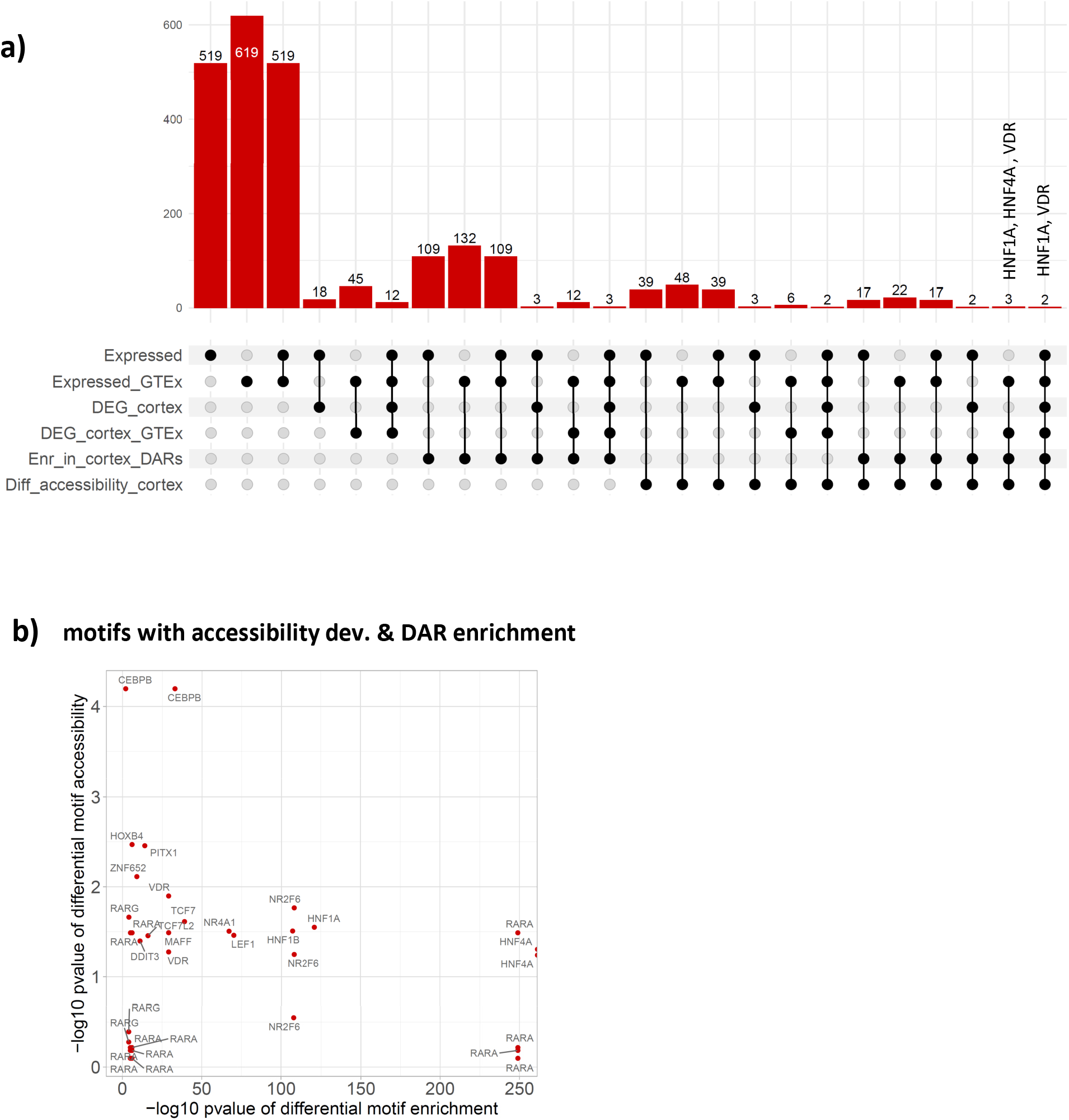
Identification of TFs driving cortical gene expression. **a)** 519 of all 661 TFs in the JASPAR 2022 CORE vertebrates database showed at least minimal expression in our an the GTEx V8 kidney samples. The upset plot shows the number of TFs which are furthermore differentially expressed in cortex in our and /or the GTEx data (DEG_cortex, DEG_cortex_GTEx), show significant motif enrichment in cortical DARs (Enr_in_cortex_DARs) and show significant medullary accessibility deviations at their motifs (Diff_accessibility_cortex). Only HNF1A and VDR were identified consistently across all analyses. **b)** Comparison of p-values from differential motif enrichment and accessibility deviations.

**Supplementary Figure 10:**
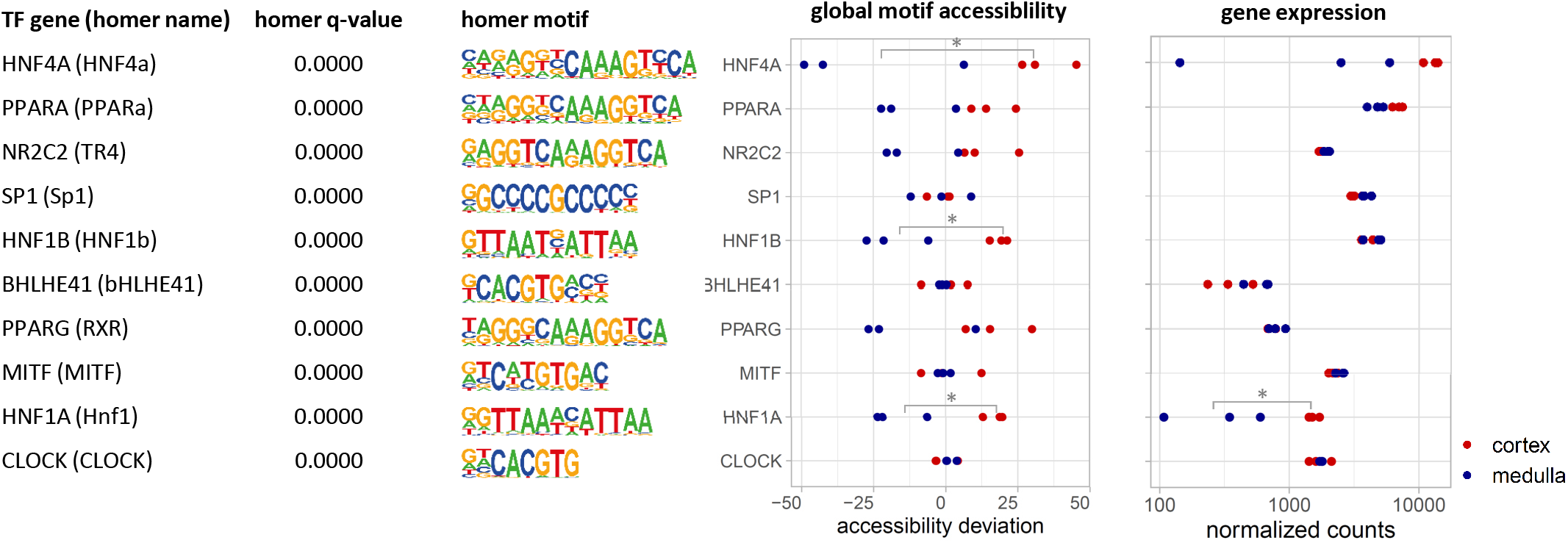
Significant TF motif enrichment in ARs within promoters of cortex DEGs. TFs with significant motif enrichment in ARs inside of medullary DEG promoters (−5000 bp – 1000 bp from TSS) computed using HOMER. The middle panel shows the accessibility deviations for these same TFs computed with chromVar for each of our cortical and medullary tissue samples. The right panel displays the TFs’ normalized expression for each sample.

**Supplementary Figure 11:**
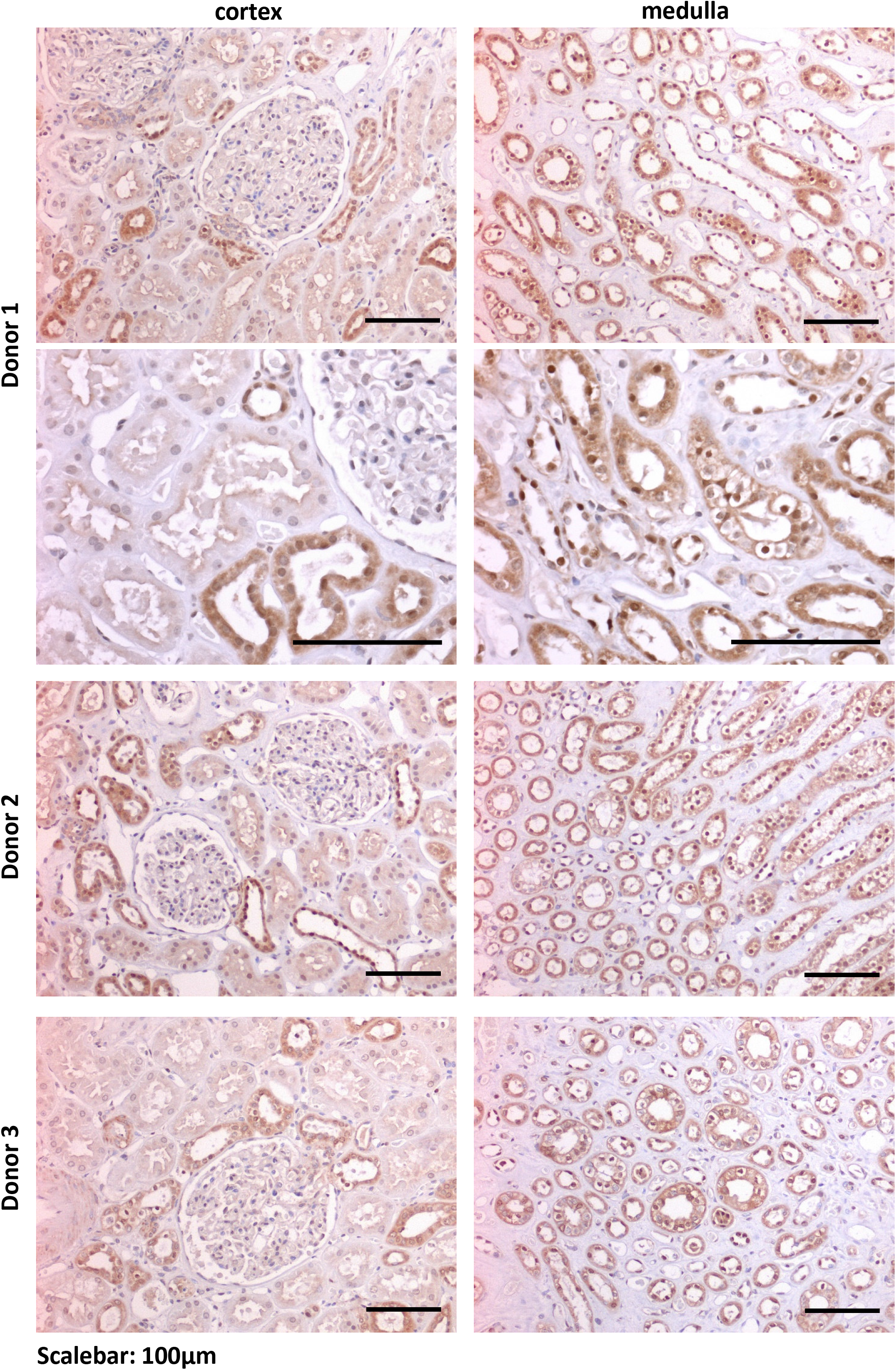
POU3F3 IHC in kidney cortex and medulla.

